# Mechanisms of insertions at a DNA double-strand break

**DOI:** 10.1101/2022.09.30.509517

**Authors:** Jaewon Min, Junfei Zhao, Jennifer Zagelbaum, Sho Takahashi, Portia Cummings, Allana Schooley, Job Dekker, Max E. Gottesman, Raul Rabadan, Jean Gautier

## Abstract

Insertions and deletions (indels) are common sources of structural variation, and insertions originating from spontaneous DNA lesions are frequent in cancer. We developed a highly sensitive assay in human cells (Indel-Seq) to monitor rearrangements at the TRIM37 acceptor locus which reports indels stemming from experimentally-induced and spontaneous genome instability. Templated insertions derive from sequences genome-wide and are enriched within 100 kb of donor regions flanking a DSB. Insertions require contact between donor and acceptor loci as well as DNA-PK catalytic activity. Notably, these templated insertions originate from actively transcribed loci, underscoring transcription as a critical source of spontaneous genome instability. Transcription-coupled insertions involve a DNA/RNA hybrid intermediate and are stimulated by DNA end-processing. Using engineered Cas9 breaks, we establish that ssDNA overhangs at the acceptor site greatly stimulate insertions. Indel-Seq revels that insertions are generated via at least three distinct pathways. Our studies indicate that insertions result from movement and subsequent contact between acceptor and donor loci followed invasion or annealing, then by non-homologous end-joining at the acceptor site.

## Introduction

Genomic insertions are rearrangements with one or several donor sequences inserted in a non-adjacent locus within the same chromosome or into a non-homologous chromosome. These rearrangements could represent up to 2% of germline copy number variations (CNV) and are frequently associated with developmental abnormalities (Carvalho and Lupski, 2016; Gu et al., 2016). Insertions and deletions are hallmarks of cancer genomes (Alexandrov et al., 2020; Cortes-Ciriano et al., 2020; Li et al., 2020). Insertions are thought to originate from mis-repair of DNA lesions, in particular DSBs. However, little is known about the underlying mechanisms responsible for these rearrangements. It has been proposed that insertions can arise from at least two distinct classes of mechanisms: cut-and-paste like events (Yu et al., 2018) or MMBIR-related processes (Gu et al., 2016; Osia et al., 2021).

If mis-repaired, DNA double strand breaks (DSBs) can lead to genome rearrangements, which are hallmarks of cancer. DSBs are predominantly repaired by nonhomologous end joining (NHEJ) in G1 and G2 phases, whereas they are repaired by homology-directed repair (HDR) in S and G2 phases by using sister chromatid as a template (Symington and Gautier, 2011). The choice between NHEJ and HDR is primarily determined by DNA end resection, a nucleolytic process in which DSB ends are converted into 3’ single-stranded DNA (ssDNA) overhangs. These ssDNA overhangs form RAD51 nucleoprotein filaments to invade into the sister chromatids for subsequent steps of HDR. Resected DNA ends can be processed by fill-in or endonucleolytic cleavage yielding blunt ends compatible with NHEJ (He and Chowdhury, 2021; Mirman et al., 2018; Zhao et al., 2021). Although these ‘post-resection’ steps potentially result in errors during NHEJ, their mutagenic consequences, such as deletion or insertion of ssDNA overhang fragments, have not been established.

DNA end resection is coupled with transcription at DSBs (Crossley et al., 2019; Marnef and Legube, 2021). In addition, DSBs occurring at actively transcribed loci induce the formation of DNA/RNA hybrid intermediates at ssDNA overhangs (Bader and Bushell, 2020). It is not clear how DNA/RNA hybrid intermediates are formed, nonetheless they are required for HDR by facilitating DNA end resection processes. Moreover, DNA/RNA hybrids might protect ssDNA overhangs from nucleolytic cleavage and deletion, which could be an unanticipated source of genome rearrangements during DSB repair (Liu et al., 2021).

DNA end resection also promotes clustering of DSBs into HDR domains, which ultimately facilitates HDR (Schrank and Gautier, 2019). Moreover, DSB clustering is in part driven by the nuclear WASP-ARP2/3-actin pathway, regulating DSB mobility which is essential for HDR (Schrank et al., 2018). But, bringing DSBs in close proximity also facilitates pathogenic translocations between distant DSBs (Zagelbaum et al., 2022). However, it is not known whether actin-dependent formation of HDR domains facilitates other genome rearrangements.

Here, we describe multiple mechanisms that generate insertions at a DSB. We establish a high-throughput sequencing methods to identify large numbers of unique insertions (Indel-Seq). Our approach reveals that donor sequences originate from experimentally-induced and spontaneous genome instability. Insertions arise genome-wide and are enriched within 100 kb donor regions flanking a DSB. Insertions require ARP2/3-dependent clustering of DSBs into HDR domains, contacts between acceptor and donor loci as well as DNA-PK catalytic activity. Transcription and DNA/RNA hybrid intermediates facilitate insertions.

## Results

### A sensitive assay to study induced and spontaneous genome insertions

We sought to investigate the mechanisms responsible for structural variation (insertions and deletions) in the human genome in response to DSBs. We used U2OS cells expressing the AsiSI endonuclease fused to the estrogen receptor ligand binding domain (AsiSI-ER) - DIvA cells (Iacovoni et al., 2010). Upon AsiSI induction with 4-hydroxy-tamoxifen (4OHT), DIvA cells experience over 100 DSBs at known genomic loci, among which the TRIM37 locus is prone to rearrangements (Cohen et al., 2018). Using γH2AX ChIP-seq and BLESS data (Aymard et al., 2014; Clouaire et al., 2018), we ranked the most frequently cut AsiSI sites and we found that the TRIM37 locus is the second most frequently cleaved site (Table 1) and (Iannelli et al., 2017). We thus elected to identify structural variants at this locus. We chose a PCR-based approach to monitor AsiSI-dependent alterations (Fig. 1A). Genomic DNA was purified 72 h following incubation +/- 4-OHT and subjected to *in vitro* AsiSI digestion to eliminate undigested, intact loci. AsiSI-resistant loci were amplified by PCR spanning the AsiSI cut site to characterize rearrangements. We observed *de novo* generation of higher molecular weight amplicons by agarose gel electrophoresis, indicative of insertion events (Fig. 1B) as well as smaller molecular weight DNA on the bioanalyzer profile, indicative of deletions (Fig. S1A). 162 unique amplicons were further analyzed by Sanger sequencing (Fig. S1B-C). Notably, 39% of inserts originate from repetitive sequences, including telomeres, centromeres, Alu, and retrotransposon elements (Fig. S1B). Next, we developed a high-throughput targeted sequencing approach combined with a robust computational pipeline that we named Indel-Seq (Insertion and deletion sequencing). We used MiSeq (Illumina) to generate 600 bp reads, including insert sequence(s) of up to 260 bp (Fig 1A). Both Sanger sequencing and Indel-Seq indicate that the median size of inserts is 120 bp (Fig. S1C-D). Donor sequences originate from all chromosomes (Fig. S1E-F), but are more frequent for chromosome 17 where TRIM37 is located, and which is enriched in incised AsiSI sites (Fig. S1G). In keeping with Sanger sequencing, Indel-Seq identified insertions originating from repetitive sequences. 28% of insertions arose from telomeres (Fig. S1B), consistent with the documented telomere instability of U2OS cells (Lee et al., 2014). Probing TRIM37 amplicons with a telomeric probe revealed a major product ~500 bp, which corresponds to 150-160 bp of telomeric repeats (Fig. S1H).

**Figure 1.**
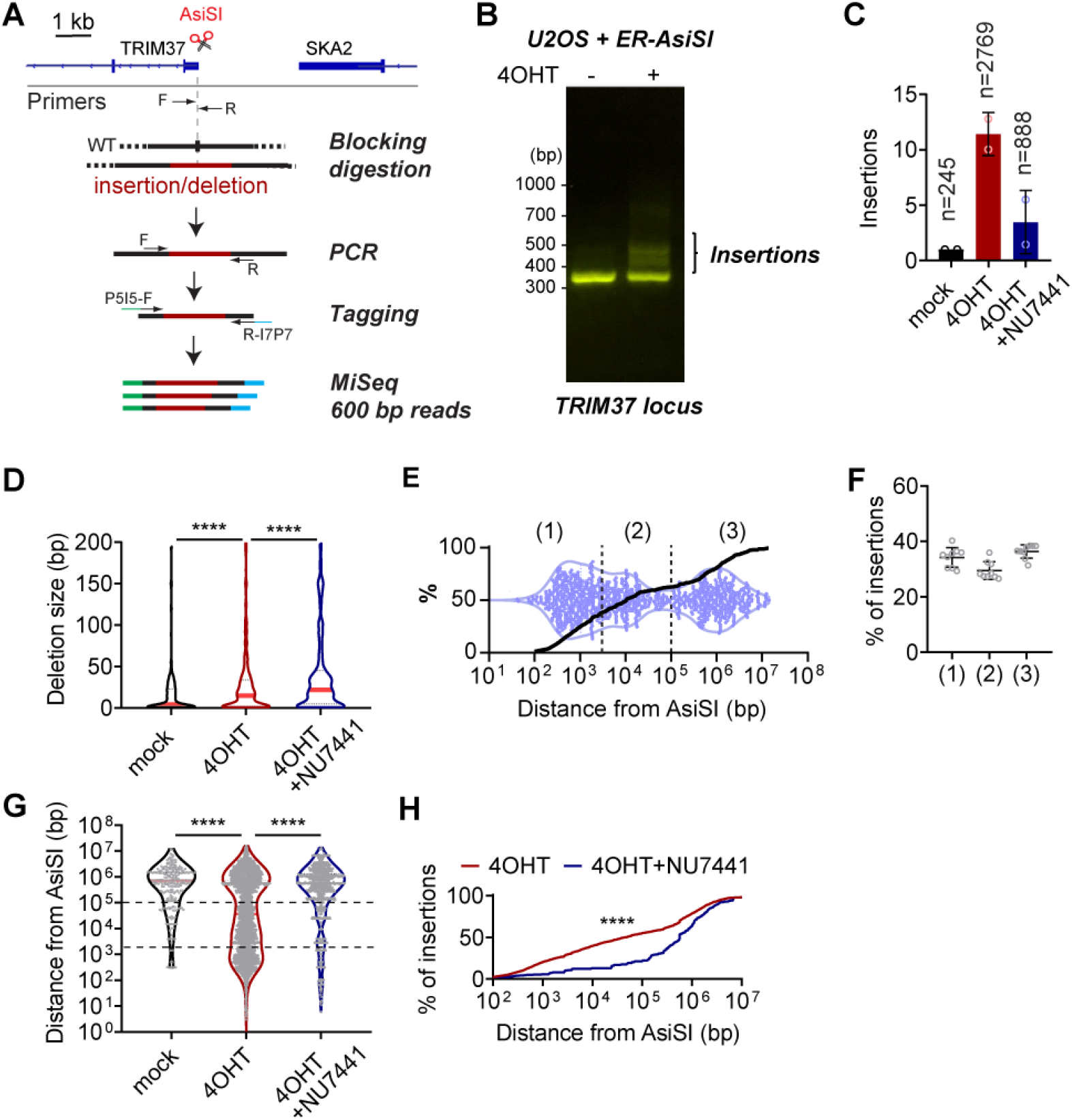
Indel-Seq assay. (A) Schematic of Indel-Seq assay to monitor insertions and deletions at a DSB. Rearrangements following AsiSI cleavage near the TRIM37 TSS are amplified and sequenced in a high-throughput manner. (B) Agarose gel electrophoresis of PCR amplicons spanning the AsiSI site at the TRIM37 locus in the presence or absence of DSBs (+/-4OHT). (C) Relative number of insertion events in control DIvA cells (mock), cells treated with 4OHT, and cells treated with 4OHT and with μM of DNA-PK inhibitor (NU7441). Columns are normalized to the frequency of insertions in control. Mean and standard deviation. (D) Deletion size shown as violin plots in control DIvA cells (mock), cells treated with 4OHT, and cells treated with 4OHT and NU7441. Red lines indicate median. *P* calculated with Student’s two-tailed t-test. (E) Violin plot (purple) and cumulative frequency of the distribution of donor sequences (black line) as a function of the distance from the nearest AsiSI site. (1) 0 – 2 kb, (2) 2 – 100 kb and (3) >100 kb from the nearest AsiSI site. (F) Frequency of donor sequences for each insertion Class in DIvA cells treated with 4OHT. The results of nine independent experiments were presented. (G) Distribution of donor sequences as a function of the distance to the nearest AsiSI site in control DIvA cells (mock), cells treated with 4OHT, and cells treated with 4OHT and NU7441. *P* calculated with Mann-Whitney test. Dashed lines delineate the 3 classes of insertions. (H) Cumulative frequency of the distribution of donor sequence in 4-OHT-treated DivA cells +/- NU7441 as a function of the distance from the nearest AsiSI site. *P* calculated with Kolmogorov-Smirnov test.

**Table 1.**
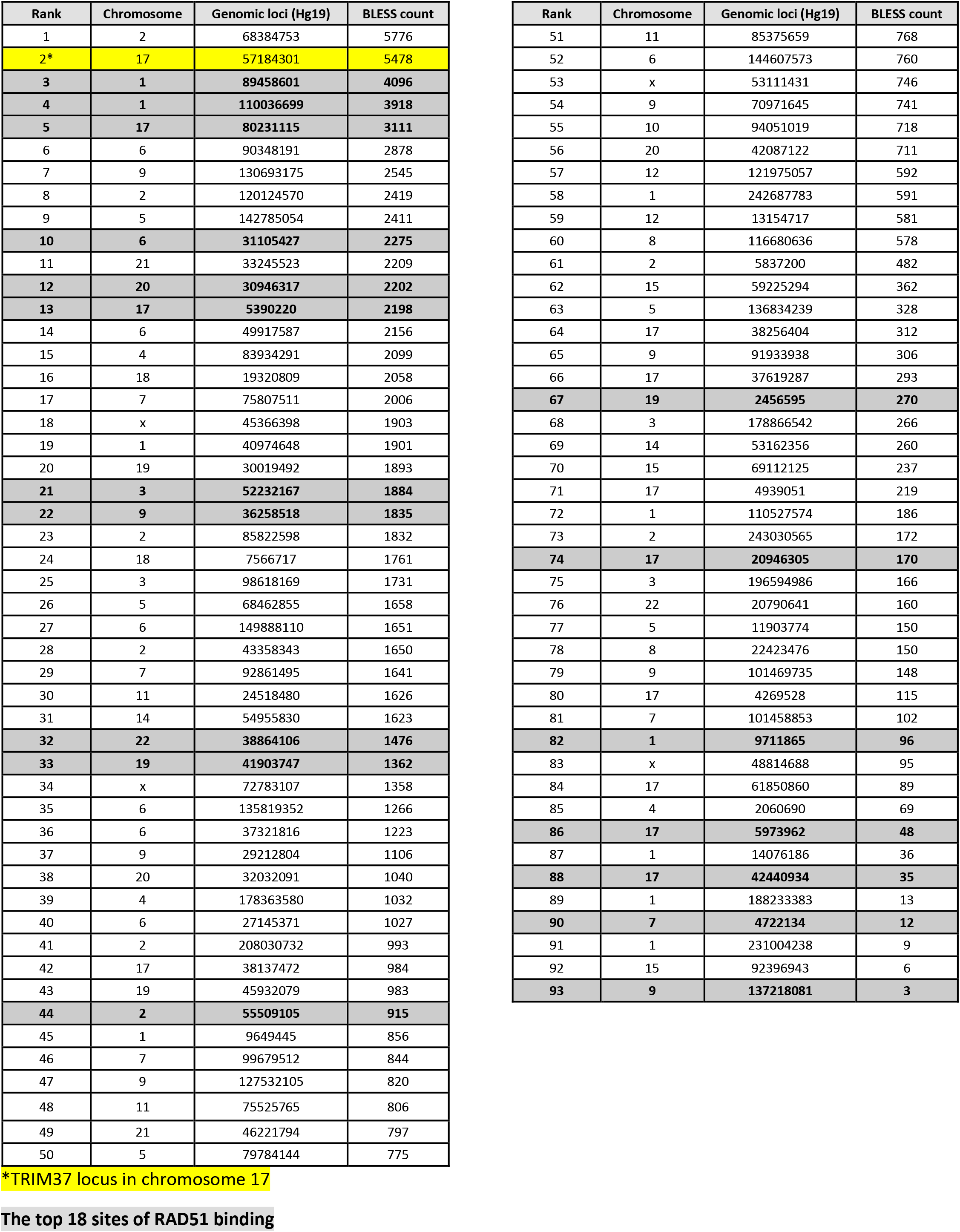
Most frequently cut AsiSI sites based on BLESS count.

| Rank | Chromosome | Genomic loci (Hg19) | BLESS count |
| --- | --- | --- | --- |
| 1 | 2 | 68384753 | 5776 |
| **2\*** | **17** | **57184301** | **5478** |
| **3** | **1** | **89458601** | **4096** |
| **4** | **1** | **110036699** | **3918** |
| **5** | **17** | **80231115** | **3111** |
| 6 | 6 | 90348191 | 2878 |
| 7 | 9 | 130693175 | 2545 |
| 8 | 2 | 120124570 | 2419 |
| 9 | 5 | 142785054 | 2411 |
| **10** | **6** | **31105427** | **2275** |
| 11 | 21 | 33245523 | 2209 |
| **12** | **20** | **30946317** | **2202** |
| **13** | **17** | **5390220** | **2198** |
| 14 | 6 | 49917587 | 2156 |
| 15 | 4 | 83934291 | 2099 |
| 16 | 18 | 19320809 | 2058 |
| 17 | 7 | 75807511 | 2006 |
| 18 | x | 45366398 | 1903 |
| 19 | 1 | 40974648 | 1901 |
| 20 | 19 | 30019492 | 1893 |
| **21** | **3** | **52232167** | **1884** |
| **22** | **9** | **36258518** | **1835** |
| 23 | 2 | 85822598 | 1832 |
| 24 | 18 | 7566717 | 1761 |
| 25 | 3 | 98618169 | 1731 |
| 26 | 5 | 68462855 | 1658 |
| 27 | 6 | 149888110 | 1651 |
| 28 | 2 | 43358343 | 1650 |
| 29 | 7 | 92861495 | 1641 |
| 30 | 11 | 24518480 | 1626 |
| 31 | 14 | 54955830 | 1623 |
| **32** | **22** | **38864106** | **1476** |
| **33** | **19** | **41903747** | **1362** |
| 34 | x | 72783107 | 1358 |
| 35 | 6 | 135819352 | 1266 |
| 36 | 6 | 37321816 | 1223 |
| 37 | 9 | 29212804 | 1106 |
| 38 | 20 | 32032091 | 1040 |
| 39 | 4 | 178363580 | 1032 |
| 40 | 6 | 27145371 | 1027 |
| 41 | 2 | 208030732 | 993 |
| 42 | 17 | 38137472 | 984 |
| 43 | 19 | 45932079 | 983 |
| **44** | **2** | **55509105** | **915** |
| 45 | 1 | 9649445 | 856 |
| 46 | 7 | 99679512 | 844 |
| 47 | 9 | 127532105 | 820 |
| 48 | 11 | 75525765 | 806 |
| 49 | 21 | 46221794 | 797 |
| 50 | 5 | 79784144 | 775 |

**\*TRIM37 locus in chromosome 17**
| Rank | Chromosome | Genomic loci (Hg19) | BLESS count |
| --- | --- | --- | --- |
| 51 | 11 | 85375659 | 768 |
| 52 | 6 | 144607573 | 760 |
| 53 | x | 53111431 | 746 |
| 54 | 9 | 70971645 | 741 |
| 55 | 10 | 94051019 | 718 |
| 56 | 20 | 42087122 | 711 |
| 57 | 12 | 121975057 | 592 |
| 58 | 1 | 242687783 | 591 |
| 59 | 12 | 13154717 | 581 |
| 60 | 8 | 116680636 | 578 |
| 61 | 2 | 5837200 | 482 |
| 62 | 15 | 59225294 | 362 |
| 63 | 5 | 136834239 | 328 |
| 64 | 17 | 38256404 | 312 |
| 65 | 9 | 91933938 | 306 |
| 66 | 17 | 37619287 | 293 |
| **67** | **19** | **2456595** | **270** |
| 68 | 3 | 178866542 | 266 |
| 69 | 14 | 53162356 | 260 |
| 70 | 15 | 69112125 | 237 |
| 71 | 17 | 4939051 | 219 |
| 72 | 1 | 110527574 | 186 |
| 73 | 2 | 243030565 | 172 |
| **74** | **17** | **20946305** | **170** |
| 75 | 3 | 196594986 | 166 |
| 76 | 22 | 20790641 | 160 |
| 77 | 5 | 11903774 | 150 |
| 78 | 8 | 22423476 | 150 |
| 79 | 9 | 101469735 | 148 |
| 80 | 17 | 4269528 | 115 |
| 81 | 7 | 101458853 | 102 |
| **82** | **1** | **9711865** | **96** |
| 83 | x | 48814688 | 95 |
| 84 | 17 | 61850860 | 89 |
| 85 | 4 | 2060690 | 69 |
| **86** | **17** | **5973962** | **48** |
| 87 | 1 | 14076186 | 36 |
| **88** | **17** | **42440934** | **35** |
| 89 | 1 | 188233383 | 13 |
| **90** | **7** | **4722134** | **12** |
| 91 | 1 | 231004238 | 9 |
| 92 | 15 | 92396943 | 6 |
| **93** | **9** | **137218081** | **3** |

We detected low levels indels at the TRIM37 locus in the absence of AsiSI induction (Fig. 1C, S1D, mock). Nevertheless, DSB induction greatly increased the number of insertions (Fig. 1C, G) and yielded larger deletions (Fig. 1D). Of note, the TRIM37 locus is amplified 1.8-fold in U2OS cells, which might facilitate detecting insertions (Cerami et al., 2012).

To probe the connection between the origin of insert sequences and loci harboring active AsiSI sites, we graphed the number of donor sequences as a function of their distance to the nearest AsiSI site, shown as violin plots or cumulative distribution (Fig. 1E). One-third of donor sequences originated from: (1) processed DSBs, immediately adjacent to AsiSI sites (0 - 2 kb); (2) one-third from a broader 100 kb region surrounding AsiSI sites, and (3) one-third from AsiSI-distal genomic loci (>100 kb) (Fig. 1F). Proximal donor sequences (1) and (2) are potentially dependent on AsiSI-induced DSBs, whereas distal insertions are presumably derived from spontaneous lesions in the genome (Fig. 1E). Donor sequences in DIvA cells that were not induced by 4OHT (mock) arise primarily from AsiSI-distal loci, further suggesting that they arise from spontaneous lesions (Fig 1E, G). Furthermore, Class (1) and (2) inserts are AsiSI-dependent (4OHT vs. mock, Fig. 1G). Class (1) might arise directly from processing of AsiSI DNA ends, which is less than 5 kb in human cells (Zhou et al., 2014).

Insertions could arise via a “cut and paste” process by direct end-joining ligation at the acceptor site. Thus, we asked if generation of indels required NHEJ. We treated cells with NU7441, a DNA-PK inhibitor to reduce non-homologous end joining (NHEJ). Inhibition of DNA-PK catalytic activity significantly decreased insertion events (Fig. 1C, G and H). In contrast, the size and numbers of deletions increased, possibly due to enhanced DNA end-resection upon inhibition of non-homologous end joining (NHEJ) (Fig. 1D) (Zhou et al., 2014). Treatment with NU7441 reduced predominantly DSB-dependent, AsiSI-proximal insertions: (1) and (2), and to a lesser extent AsiSI-distal (3) insertions (Fig. 1G-H). These data establish that Indel-Seq can readily detect experimentally induced insertions originating from the vicinity of AsiSI cleaved loci as well as insertions arising from endogenous DNA lesions. Our data indicate that deletion and insertions originating from AsiSI-proximal loci require DNA-PKcs, suggesting the involvement of NHEJ.

### DSB clustering into HDR domain facilitates single and multiple insertions

DSBs induce chromatin movement and clustering of damaged loci (Aten et al., 2004; Lisby et al., 2003), which is in part driven by the nuclear WASP-Arp2/3-Actin pathway (Caridi et al., 2018; Schrank et al., 2018). DSB clustering within HDR domains, however, can facilitate rare pathological chromosome translocations (Zagelbaum et al., 2022). We hypothesized that contact between broken chromosome ends would favor transient interactions yielding genomic insertions. We therefore assessed the impact of inhibiting Arp2/3-mediated DSB mobility on insertion events. Treatment with CK-666, a small molecule inhibitor which stabilizes the Arp2/3 complex in an inactive conformation (Nolen et al., 2009), resulted in a significant decrease in insertion events originating from AsiSI-proximal donor sequences (Fig. 2A-B, S2A-B), strongly suggesting that ARP2/3-dependent forces and subsequent DSB clustering into HDR domains could elicit contact between donor and acceptor loci and greatly facilitate genomic insertions. To assess directly the cxonnection between broken chromosome contacts and insertions, we monitored genome interaction upon DSB induction. We analyzed chromatin contacts by chromosome conformation capture (Hi-C) in DIvA cells. We monitored intra-chromosomal interactions between the acceptor site (TRIM37 – magenta vertical bar) on chromosome 17 and the rest of chromosome 17, normalized to observed/expected values (Fig. 2C, top). Strengthened differential interactions following DSBs are shown in red above the x-axis whereas blue bars below the x-axis represent weakened interactions (Fig. 2C, top). The number and location of donor sequences along chromosome 17 are shown below (Fig. 2C, bottom). The TRIM37 locus itself is excluded from the interaction analysis. These results indicate that genome interactions were significantly enhanced between TRIM37 and frequently cleaved AsiSI sites (Fig. 2C, red dash lines), consistent with our studies in mouse embryonic fibroblasts (MEFs) (Zagelbaum et al., 2022). Moreover, donor sequences originated preferentially from loci harboring enhanced interactions following DSBs (Fig 2C, bottom). This strongly suggests that insertions are mediated by contacts between the acceptor and donor loci.

**Figure 2.**
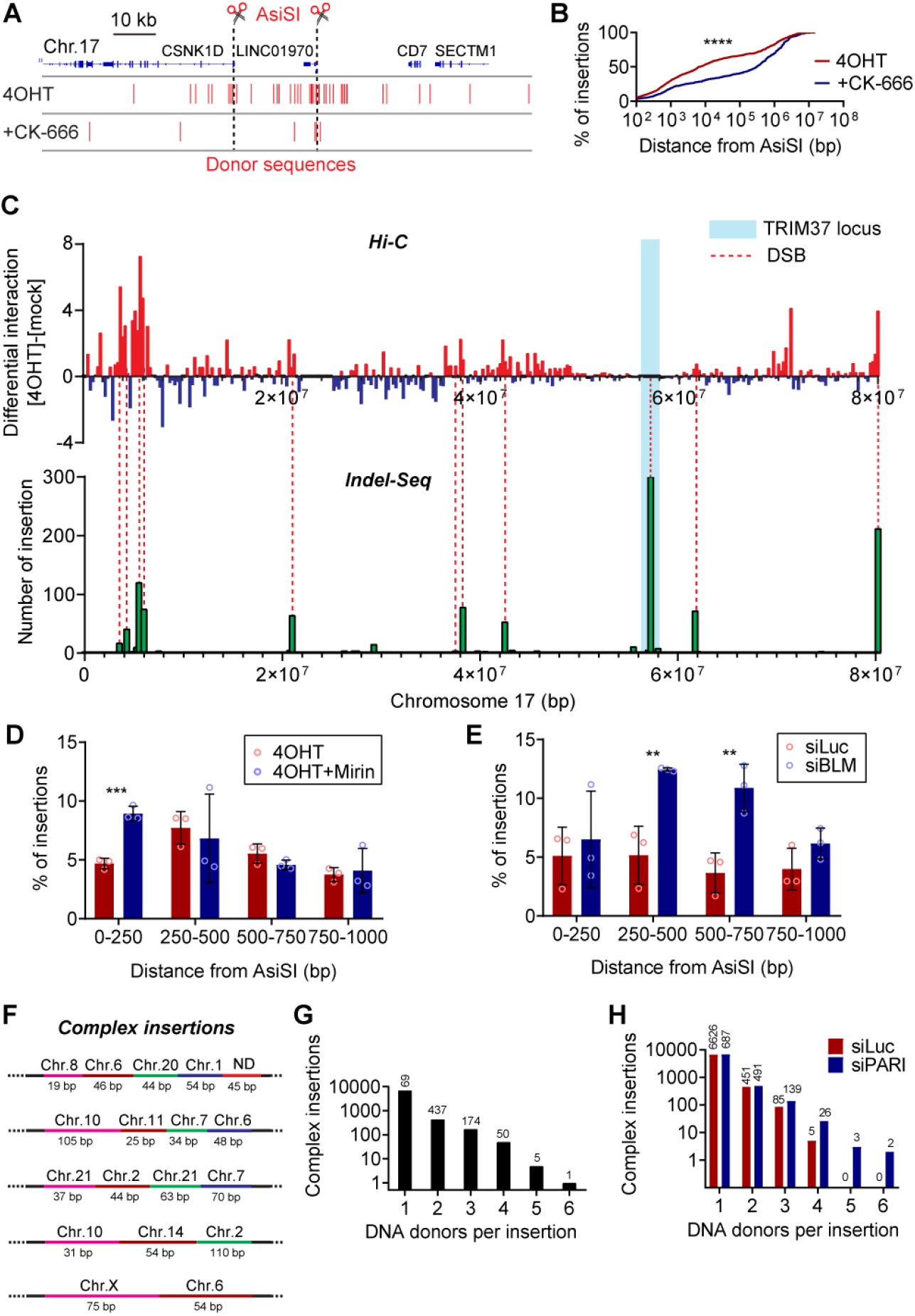
Arp2/3-mediated DSB clustering drives insertions. (A) Genome browser view of the CSNK1D locus on chromosome 17 (top) and position of donor sequences in DivA cells treated with 4OHT (middle) and CK-666 (bottom). The black dashed line represents AsiSI-induced DSB site. (B) Cumulative frequency of the distribution of donor sequences in DivA cells treated with 4OHT +/- CK-666 as a function of the distance from the nearest AsiSI site. *P* calculated by Kolmogorov-Smirnov test. (C) Top: Differential interaction plot from Hi-C data of DivA cells in the presence and absence of DSBs normalized to observed/expected between a 1 Mb region surrounding the acceptor site (TRIM37 locus, blue box) on chromosome 17 and the rest of chromosome 17. Bottom: number of donor sequences inserted at the acceptor site following DSB induction. Bin size is 250 kb. (D) Frequency of donor sequences in DIvA cells treated with 4OHT +/- Mirin (50 μM) as a function of the distance from the nearest AsiSI site. The histogram bin size is 250 bp. *P* calculated with Student’s two-tailed t-test. (E) Frequency of donor sequences in BLM-depleted and control cells as a function of the distance from the nearest AsiSI site. The histogram bin size is 250 bp. *P* calculated by Student’s two-tailed t-test. (F) Five examples of complex insertions at acceptor sites. (G) Number of donor sequences per insertion sites in 4-OHT treated DivA cells. (H) Number of donor sequences per insertion sites in PARI-depleted (siPARI) and control (siLuc) cells.

One third of insertions arise from within 2 kb of cleaved AsiSI sites (Fig. 1F) and therefore could be templated from resected DNA ends (Zhou et al., 2014). Moreover, DNA end-resection is coupled to DSB mobility (Schrank et al., 2018) and clustering (Aymard et al., 2017) prompting us to assess the role of end-processing in mediating insertions. For short-range resection, we evaluated the impact of mirin, a small molecule inhibitor of MRE11 endo- and exonuclease activities (Dupre et al., 2008), on genomic insertions. MRE11 inhibition by mirin led to a decrease in the number of insertions (Fig. S2C), indicating that DNA end-resection facilitates insertion events. To assess the contribution of long-range resection on the distribution of donor sequences, we down-regulated BLM (Fig. S2E), which performs long-range resection as a complex with DNA2 in parallel with the EXO1 nuclease (Nimonkar et al., 2011). Notably, treatment of DIvA cells with 50 μM mirin increased the fraction of the most AsiSI-proximal donor sequences originating from within 250 bp of DSBs (Fig. 2D, Fig. S2D), whereas BLM downregulation increased the frequency of donor sequences originating further away, albeit still within 1 kb of DSBs (Fig. S2F). Thus, both BLM and Mre11 inhibition favored donor sequences originating close to DSBs (Fig. 2D-E, S2D, F). These data suggest that ssDNA generated at processed DSBs could be a template for donor sequences. Indeed, the length of resection tracks: WT > BLMi > Mre11i, correlates with the position of donor sequences vis-à-vis AsiSI sites.

RAD51 regulates DSB mobility downstream of DNA end-resection (Dion et al., 2012; Mine-Hattab and Rothstein, 2012). We assessed the extent of chromatin-bound RAD51 at cleaved AsiSI loci using RAD51 ChIP-Seq data (Aymard et al., 2017) and BLESS data from Table 1 (Clouaire et al., 2018). We ranked AsiSI sites from Table 1 for chromatin-bound Rad51 and found the 20% most enriched loci for Rad51 contributed 50% of donor sequences (top 18 Rad51-enriched DSBs, Fig. S2G). Furthermore, donor sequences in these top18 cleaved sites originated further away from the AsiSI sites (Fig. S2H). Thus, donor sequences preferentially originate from RAD51-enriched DSBs, further implicating HDR recombination mechanisms.

Indel-Seq revealed that 8.7% of insertions originated from multiple genomic loci on different chromosomes (Fig. 2F). These complex insertions events involve up to 6 donors (Fig. 2F, 2G). We hypothesized that complex insertions result from multiple, successive invasion or annealing events, possibly involving homologous recombination.

To test this hypothesis, we altered the levels of chromatin-bound RAD51. Human PARI, similar to yeast Srs2, antagonizes HDR by disrupting the formation of RAD51 filaments (Moldovan et al., 2012). Loss of PARI leads to hyperrecombination phenotypes, including increased RAD51 foci and sister chromatid exchanges. PARI downregulation in DIvA cells increased the frequency of complex insertions as well as the number of donor sequences per insertion (Fig. 2H). This further suggests that single and multiple DSB-dependent insertions are mediated by RAD51 recruitment and DSB clustering.

Collectively, our data indicate that genome insertions arise following clustering of DNA DSBs into HDR domains as well as a subsequent RAD51-dependent recombination step. This implies a more complex mechanism beyond a simple “cut and paste”, ligation-based process.

### Genome insertions are coupled with RNA synthesis at DSBs

The most frequently cut AsiSI sites are located within accessible chromatin regions proximal to transcription start sites (TSS) (Fig S3A). In addition, more than 70% of donor sequences originated from promoters and transcribed gene body regions (Fig. 3A. Fig. S3B-C). Notably, insertions originating from promoter loci were within 1 kb of cleaved AsiSI sites whereas those originating from gene bodies were further away (Fig. 3A). We therefore hypothesized that the mechanism of insertion was coupled to transcription and examined AsiSI-induced DSBs located near a TSS. Donor sequences are spread around the AsiSI sites and span the TSS, however, insertions preferentially originate from the transcribed side (Transcription +, Fig. 3B-C). In contrast, donor sequences originating from AsiSI-cleaved DSBs within a gene body or distal intergenic regions did not show any bias and were equally distributed on each side of the AsiSI sites (Fig. S3D-E). Donor sequences originating from the transcribed side of a DSB were located farther from the AsiSI site compared to donor sequences originating from the non-transcribed side (Fig. S3F). These data suggest that active transcription facilitates the generation of donor sequences.

**Figure 3.**
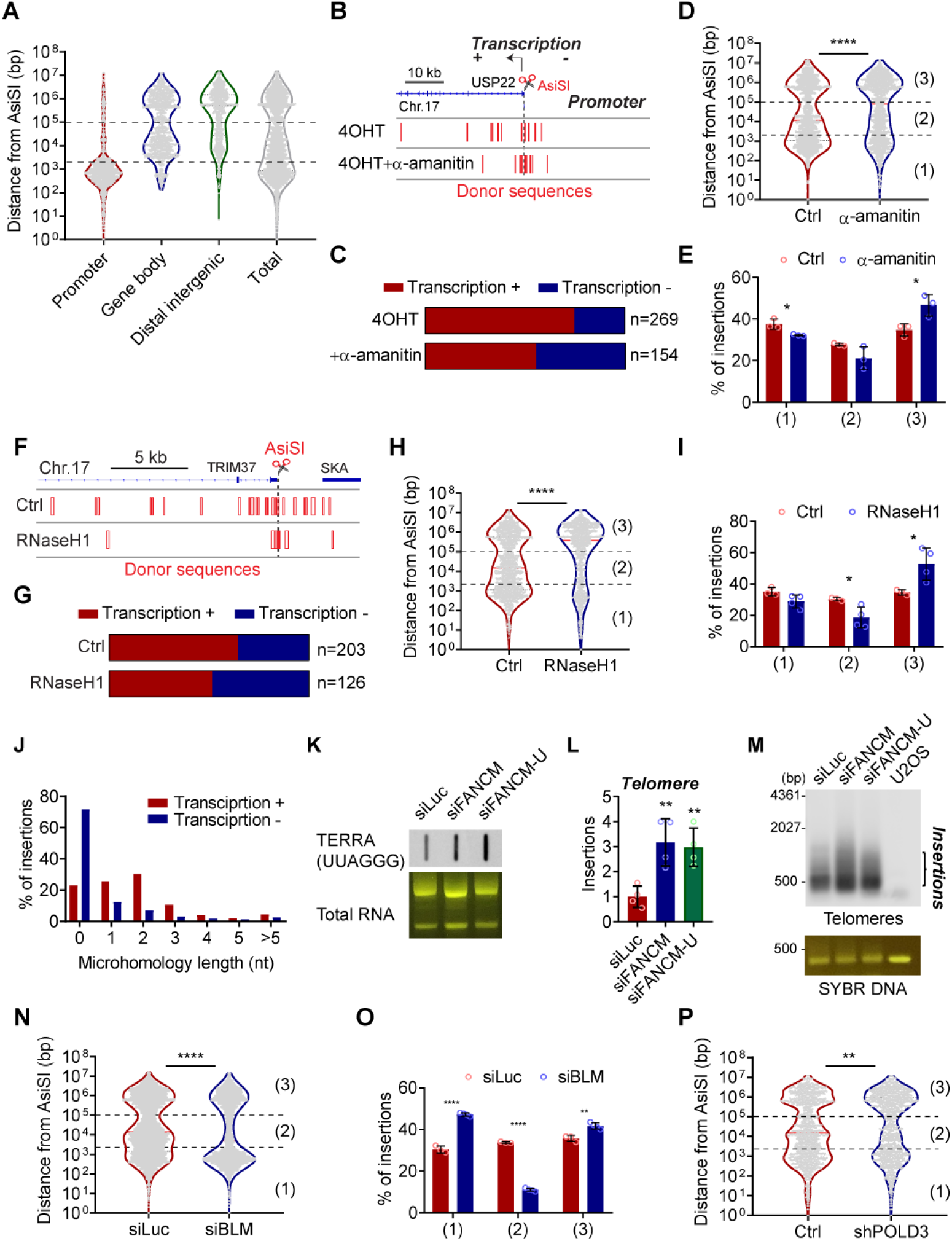
Role of transcription. (A) Distance from the nearest AsiSI site for donor sequences derived from promoter, gene body and distal intergenic regions. *P* calculated with Mann-Whitney test. (B) Genome browser view of the USP22 locus on chromosome 17 (top) the TSS is near an AsiSI site and the direction of transcription is indicated by an arrow. Donor sequences in 4-OHT treated DivA cells (middle) and cells treated with 1 μg/ml α-amanitin (bottom). The black dashed line represents AsiSI-induced DSB site. (C) Fraction of donor sequences arising from transcribed (Red, Transcription +) or un-transcribed (Blue, Transcription -) sides at loci for which cleaved DSBs are nearby TSS in DivA cells treated with 4OHT +/- α-amanitin. (D) Distribution of donor sequences as a function of the distance to the nearest AsiSI site in DivA cells treated with 4OHT +/- α-amanitin. Bar graphs represent median. (E) Frequency of sequences derived from each donot class ((1) 0 – 2 kb, (2) 2 – 100 kb and (3) >100 kb from the nearest AsiSI site). in DivA cells treated with 4OHT +/- α- amanitin. (1) 0 – 2 kb, (2) 2 – 100 kb and (3) >100 kb from the nearest AsiSI site. (F) Genome browser view of the TRIM37 locus (top). Distribution of donor sequence in control (middle) or RnaseH1 cDNA overexpressing Diva cells (bottom). The black dashed line represents AsiSI-induced DSB site. (G) Fraction of donor sequences arising from transcribed (Red, Transcription +) or un-transcribed (Blue, Transcription -) sides at loci for which cleaved DSBs are nearby TSS in control and RnaseH1 cDNA overexpressing DivA cells. (H) Distribution of donor sequences as a function of the distance to the nearest AsiSI site in control and RnaseH1 cDNA overexpressing DivA cells. (I) Frequency of donor sequences that derived from each donor class in control and RnaseH1 cDNA overexpressing DivA cells. (J) Frequency of insertion junctions in transcribed (Red, Transcription +) or un-transcribed (Blue, Transcription -) direction as a function of the length of microhomology (nt, nucleotide) (K) TERRA (UUAGGG) Northern blot (top) and total RNA (bottom) from control (siLuc) and FANCM depleted cells. siFANCM-U targets the 3’ UTR. (L) Relative number of telomere insertion events in control (siLuc) and FANCM depleted DIvA cells. Insertions are normalized to siLuc control. Mean and standard deviation. (M) Telomere (TTAGGG) Southern blot (top) and SYBR DNA staining (bottom) of amplicons spanning the AsiSI site at the TRIM37 locus in control (siLuc) DivA and FANCM depleted cells. (N) Distribution of donor sequences as a function of the distance to the nearest AsiSI site in control (siLuc) and BLM-depleted cells. (O) Frequency of donor sequences from each class in control (siLuc) and BLM-depleted cells. (P) Distribution of donor sequences as a function of the distance to the nearest AsiSI site in control and POLD3-depleted cells.

To assess directly whether transcription drives insertions, we monitored insertion events in cells treated with low doses of alpha-amanitin to partially inhibit RNA polymerase II and III (RNAPII and RNAPIII) - mediated transcription (Lindell et al., 1970; Weinmann and Roeder, 1974). We found that alpha-amanitin decreased the frequency of insertions; preferentially those originating from promoter-proximal DSBs: (1) and (2) (Fig. 3D-E, S3G). Furthermore, donor sequences in alpha-amanitin treated cells were no longer preferentially derived from the transcribed side (Fig. 3B-C).

RNA synthesis at DSBs yields DNA/RNA hybrids (R-loops) which enable DNA end-resection (Marnef and Legube, 2021). Specifically, the TRIM37 locus accumulates R-loops in response to DSB formation (Fig. S3H, bottom; S9.6 ChIP-Seq) (Cohen et al., 2018) and RNA-DNA hybrids correlate positively with DSB frequency, (BLESS signal) (Fig. S3I). Transcription-coupled RNA-DNA hybrids at donor sites might facilitate the generation of insertions by providing an intermediate for strand invasion and DNA synthesis. To determine if a DNA-RNA intermediate served as a source of insertions, we overexpressed RNaseH1 to resolve R-loops. RNA-DNA hybrid disruption by RNaseH1 suppressed the bias of donor sequences that originate preferentially from the direction of transcription, suggesting that transcription-dependent RNA-DNA hybrids are an intermediate for insertions (Fig. 3F-G). RNaseH1 overexpression resulted in a significant switch in the distribution of donor sequences, reducing primarily categories (1) and (2) within 100 kb from AsiSI sites (Fig. 3H-I, S3J). This decrease was particularly marked for donor sequences originating from promoter-proximal DSBs (Fig. S3K). Next, we examined the nucleotide sequences at insertions junctions and found a fraction of insertions harboring 2-3 nt. microhomologies (MHs). Notably, these MHs showed a strong transcription bias (Fig. 3J). Insertions arising from the direction of transcription harbor MHs, whereas insertions arising from the opposite direction do not. In addition, 59% of insertions have one blunt end and one end with MHs, supporting the idea that insertion events are initiated by microhomology-mediated strand invasion and completed by NHEJ.

We reasoned that if transcription facilitates insertions, increasing transcription locally should yield more insertions. Transcription of TERRA at telomeres results in R-loop accumulation and subsequent BLM- and POLD3-dependent break-induced replication (BIR) yielding genome instability (Arora et al., 2014; Dilley et al., 2016; Min et al., 2019; Sobinoff et al., 2017). TERRA transcription is limited by FANCM activity and down-regulation of FANCM increases transcription at telomeres (Pan et al., 2019; Silva et al., 2019). We confirmed this finding using 2 distinct siRNA targeting FANCM (Fig. 3K). Strikingly, increased TERRA transcription resulted in a significant increase of telomeric insertions as measured by Indel-Seq (Fig. 3L), a finding confirmed by Southern blot (Fig. 3M). Next, we downregulated BLM (Fig. S2E) and POLD3, the non-essential subunit of DNA polymerase delta (Fig. S3L), using siRNAs. Down-regulation of both BLM and POLD3 resulted in a dramatic shift in the distribution of donor sequences, those originating 2-100 kb from AsiSI sites (2) being the most decreased (Fig. 3M-P), consistent with RNase H overexpression (Fig. 3I). Telomere insertions were decreased in BLM depleted conditions (Fig. S3M).

Collectively, our data indicate that transcription-coupled formation of RNA-DNA hybrids directly facilitates genomic insertions, a process mediated in part by BIR.

### Single-strand DNA overhangs facilitate insertions

Next, we asked whether the nature of the DSB at the acceptor site influenced genome insertions and deletions. WT-Cas9 and mutant Cas9 nickases were used to generate blunt DSBs or breaks harboring 5’ or 3’, 46 bp ssDNA overhangs (Vriend et al., 2016). We designed two gRNAs targeting the acceptor TRIM37 locus near the original AsiSI site used in previous experiments, yielding nicks on opposite DNA strands (paired nicks). Cas9 D10A nickase, harboring a mutation in its RuvC domain, only cleaves the complimentary strand to the gRNA and creates 5’ overhangs, whereas Cas9 H840A nickase, harboring a mutation in its HNH domain, only cleaves the non-complimentary strand to the gRNA and creates 3’ overhangs. Cas9 WT generates blunt ends (Fig. 4A).

**Figure 4.**
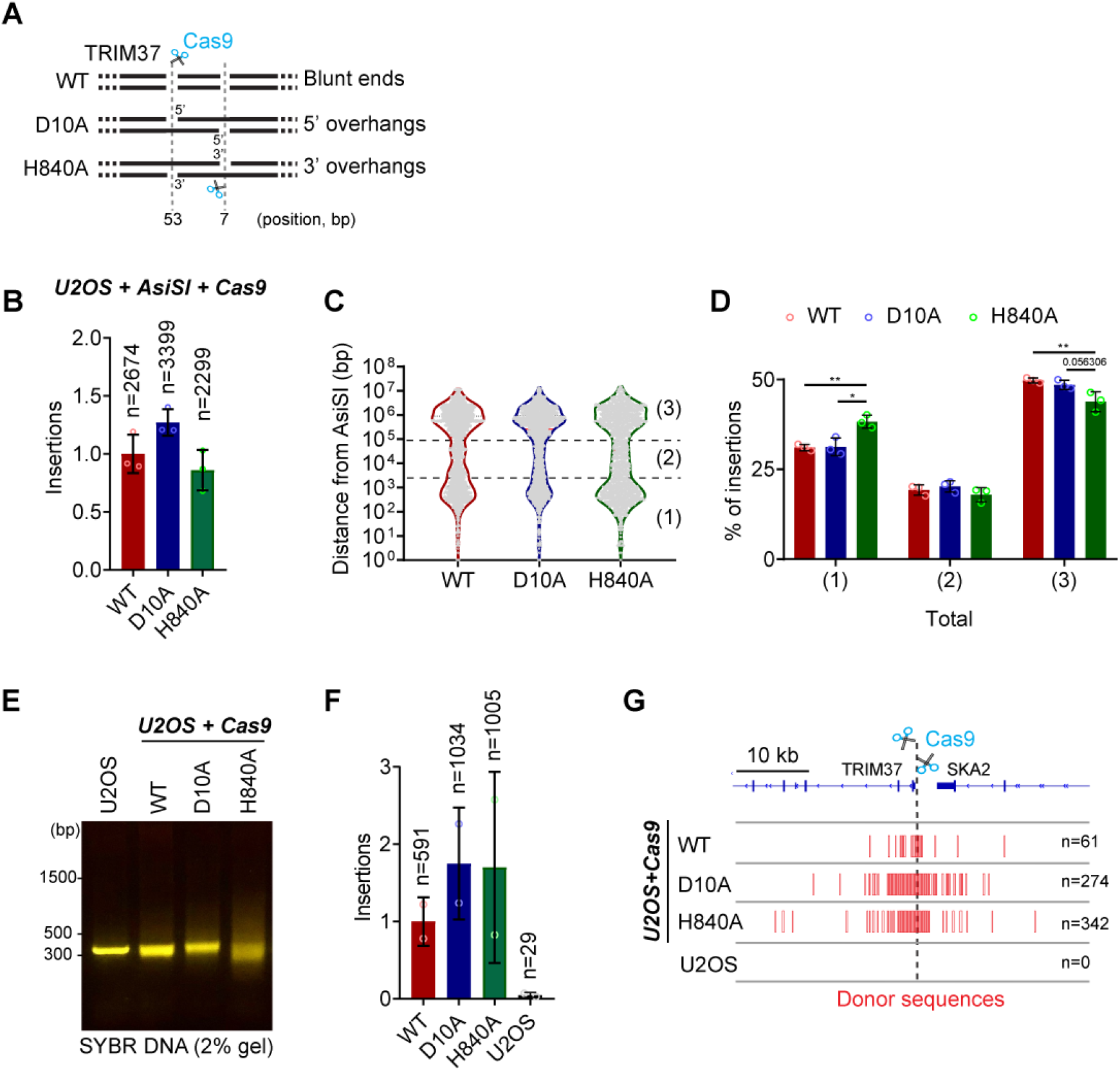
Insertions at Cas9 breaks. (A) Scheme of the paired nicks system to generate 5’ or 3’ overhangs with Cas9 nickases (D10A, H840A). (B) Relative number of insertion events in DivA cells expressing AsiSI and WT or nickases Cas9 and gRNAs targeting TRIM37 locus. Insertion frequencies are normalized to DivA cells expressing WT Cas9. Mean and standard deviation. (C) Distribution of donor sequences as a function of the distance to the nearest AsiSI site WT and nickases Cas9 expressing in DivA cells. (D) Frequency of donor sequences from each class in WT and nickases Cas9 expressing DivA cells. (E) Staining of amplicons spanning the AsiSI site at the TRIM37 locus in U2OS cells expressing WT and nickases Cas9 and gRNAs targeting TRIM37 locus. (F) Relative number of insertion events in U2OS cells expressing WT and nickases Cas9 and gRNAs targeting TRIM37 locus. Insertion frequencies are normalized to U2OS cells expressing WT Cas9. Mean and standard deviation. (G) Genome browser view of the TRIM37 locus (top). Donor sequence distribution in U2OS cells expressing WT and nickases Cas9 and gRNAs targeting TRIM37 locus (bottom).

We introduced lentiviral Cas9 WT and the mutant nickase constructs together with the gRNAs in U2OS cells that express AsiSI-ER (DiVa cells). Cas9 constructs were expressed for 24 hrs to trigger a DSB at the acceptor TRIM37 site prior to inducing AsiSI breaks genome-wide. All three types of DSBs yield similar number of insertions (Fig. 4B), however, the distribution of donor sequences genome-wide (Fig. 4C) shows a significant increase in the most AsiSI-proximal, Class (1) when the break was induced by H840A nickases (Fig. 4D, S4A). This indicates that a 3’ ssDNA overhang at the acceptor locus favors resection-dependent insertions from cleaved AsiSI sites, possibly by providing an annealing template that mimics a resected DNA end.

Next, we introduced lentiviral Cas9 WT and the mutant nickase constructs together with the gRNAs in WT U2OS cells that do not express AsiSI-ER. We found that cells expressing Cas9 nickase variants exhibited a significant increase in the average size of deletions as compared to cells expressing WT Cas9 (Fig. S4B). Median deletion sizes were 52 bp, 71 bp and 89 bp for WT, D10A and H840A, respectively (Fig. S4B). Notably, cells expressing Cas9 nickase mutants also generated substantially more insertion events, including complex insertions, compared to cells expressing Cas9 WT (Fig. 4E-F, S4C). This indicates that both 3’ and 5’ ssDNA overhangs are processed to accept donor DNA more efficiently, yielding rearrangements at the acceptor DSB. Furthermore, 26% of all donor sequences originated from the TRIM37 locus itself with a higher fraction from the locus in the nickase mutants (Fig. 4G, Table 2). 72% of insertions did not originate from the TRIM37 locus in WT-U2OS cells and likely arise from spontaneous lesions in the genome, equivalent to AsiSI distal insertions in AsiSI-ER U2OS cells (Fig. S4D, Table 2). Additionally, Cas9 breaks also generated telomere insertions, which were particularly abundant in D10A nickase 5’ ssDNA overhangs (Fig. S4E-F). Notably, 5’ ssDNA is involved in BIR processes at telomeres (Oganesian and Karlseder, 2011).

**Table 2.** The distribution of donor sequences from U2OS+Cas9 samples.

| Genomic loci | *U2OS+Cas9* |  |  |  |  |  |  |  |
| --- | --- | --- | --- | --- | --- | --- | --- | --- |
|  | WT |  | D10A |  | H840A |  | Total |  |
| Chr 17: 0-2kb from Cas9 break(s) | 51 | 9.8% | 199 | 19.2% | 289 | 28.7% | 539 | 20.9% |
| Chr 17: 2-100kb from Cas9 break(s) | 16 | 3% | 80 | 7.7% | 65 | 6.4% | 161 | 6.2% |
| Rest of chr17 | 35 | 6.7% | 61 | 5.9% | 52 | 5.2% | 148 | 5.8% |
| Rest of the genome | 419 | 80% | 694 | 67.2% | 599 | 59.6% | 1712 | 66.8% |

### Transcription associated insertions are present in cancer genomes

We reasoned that if transcription and RNA-DNA hybrids facilitates insertions, we should observe scars of transcription-associated mutations in cancer genomes. Thus, we analyzed insertions, deletions, and nucleotide variants within 100 kb regions flanking transcription start sites. We assessed mutation enrichment on the transcribed (+) or non-transcribed side (-) (transcription-associated bias) as a function of gene expression (Fig. S5A). The analysis of the Pan-Cancer Analysis of Whole Genomes (PCAWG) showed that several tumor types, including Diffuse Large B-Cell Lymphoma (DLBCL) display a positive correlation between transcription and mutagenesis (Fig S5A-B). Next, we monitored transcription associated insertions and deletions only and found that DLBCL displayed the strongest positive correlation (r=0.47, p=3.8e-05) (Fig. 5A). Of note, known lymphoma tumor suppressors (BCL6) are among the genes showing the strongest association between transcription and the presence of INDELs (Fig. 5B).

**Figure 5.**
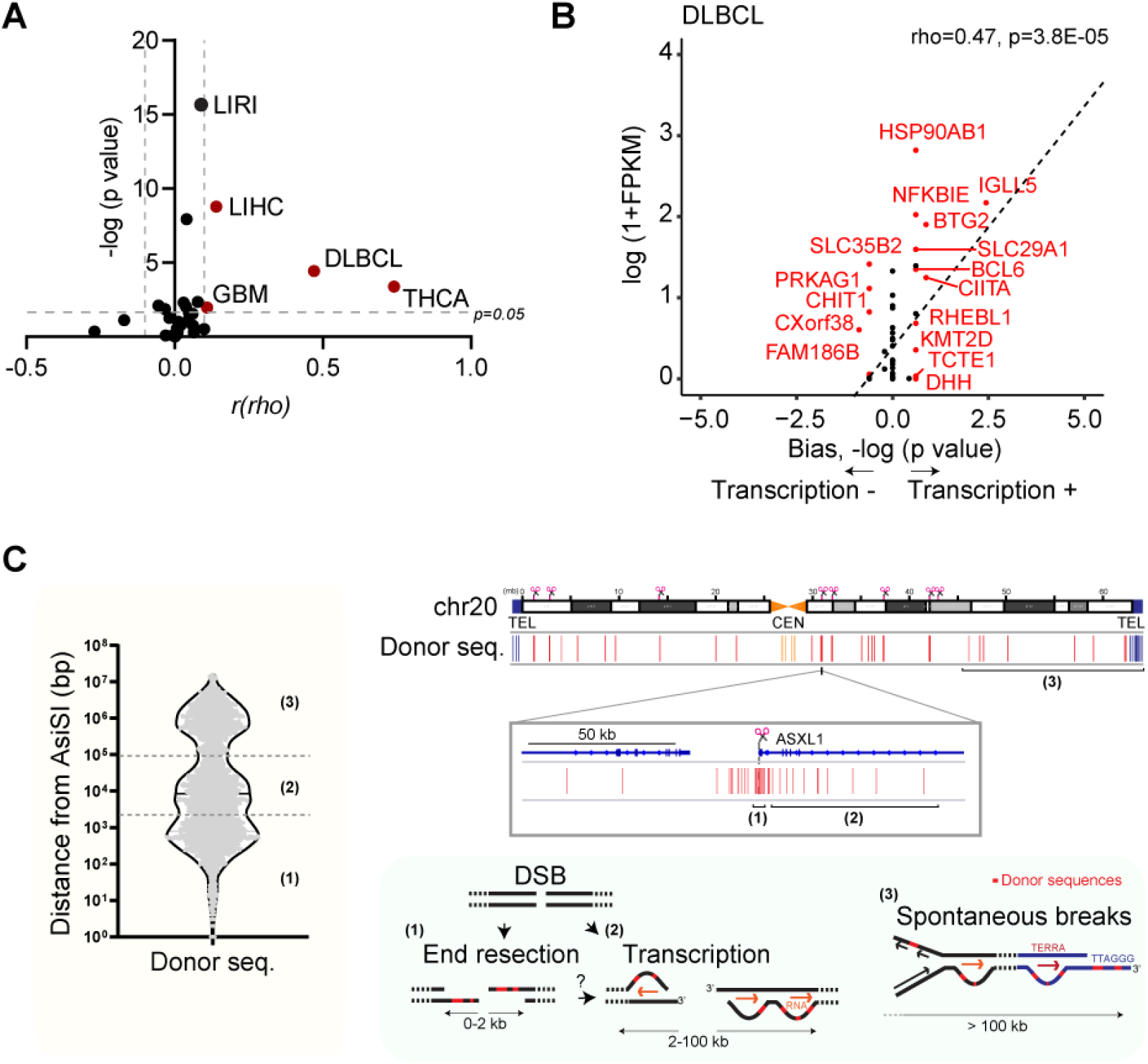
Model: origins of templated insertions. (A) Analysis of transcription-associated indel bias in 29 tumor types. x-axis: r (rho), correlation between transcription-associated bias and gene expression levels, y-axis: negative log p value. r = ± 0.1, p=0.05: cut-off. (B) The analysis of transcription-associated bias in indels in the diffuse large B cell lymphoma (DLBCL) (C) Model: Donor sequences yielding insertions fall into three classes according to their position relative to cleaved AsiSI sites. Left: genome-wide distribution of donor sequences relative to the nearest AsiSI sites. (1) 0 – 2 kb, (2) 2 – 100 kb and (3) >100 kb from the nearest site. Right, top: distribution of donor sequences along chromosome 20, red bars. Please note that one red bar represents multiple donors. All cleaved AsiSI are indicated on top. (3) below points to an AsiSI-free region of chromosome 20 with distal donor sequences. Right, middle: Distribution of donor sequences spanning a cleaved AsiSI site at the ASXL1 locus. (1) donors immediately adjacent to the AsiSI site within physiological end-resection range; (2) donors spanning a 100 kb region around the AsiSI site. Right, bottom: proposed origins of donor sequences, indicated in red. (1) annealing within ssDNA overhangs of resected DSBs, (2) invasion of displaced DNA strand at DSB- and transcription-dependent R-loops and (3) spontaneous lesions (see text) including transcription at telomeres.

## Discussion

### Indel-Seq: a sensitive and versatile method to study insertions and deletions

We developed Indel-Seq, a high-throughput sequencing method that uncovers three classes of insertions based on their genomic origins in relation to experimentally induced DSBs. These three classes reflect multiple mechanisms of insertions at a DNA double-strand breaks (Figs 1 and 5). Class 1 and 2 insertions arise proximal to AsiSI breaks, originating from resected DNA ends and actively transcribing loci surrounding AsiSI breaks, respectively. Class 3 insertions arise distal from AsiSI breaks as the consequence of physiological, endogenous lesions.

The acceptor loci can be generated by a restriction endonuclease (AsiSI) or by Cas9-dependent cleavage. In AsiSI-expressing cells undergoing multiple DSBs (Iacovoni et al., 2010; Zagelbaum et al., 2022), we propose that inserts originating > 100kb from cleaved AsiSI sites (Class 3) arise from spontaneous DNA lesions, including telomere insertions. Notably, more than half of these AsiSI-distal donors are located > 1mb away from these sites. In Cas9-expressing cells, in which a single lesion is induced at the TRIM37 locus on Chr 17, donor sequences originating from all other intact chromosomes arise from spontaneous damage. We propose that these lesions are the consequence of physiological DNA transactions: transcription and replication, associated with genome instability. Our data confirmed that Cas9 lesions can be frequently mis-repaired yielding on-target deletions and insertions (Nambiar et al., 2022). For Class 3 insertions, we document in depth the example of transcription at telomeres in U2OS, with inherent telomeres fragility. Unlike Class 1 and 2, most Class 3 insertions are not clustered in the genome and the precise mechanism yielding a specific Class 3 insertion is difficult to evaluate.

### Genome insertion require contacts between acceptor and donor sites

Study of programmed chromosome rearrangements in immune cells cemented the idea that a major step towards genome rearrangements is the direct ligation of two DNA fragments. Following RAG 1/2 induced DSBs and deletion of unused DNA sequences, rearranged fragments are ligated by classical NHEJ (Alt et al., 2013; Roth, 2003). This concept has guided the hypothesis to explain the origin of genomic insertions as a “cut and paste” mechanism in which a free, extrachromosomal piece of DNA is ligated directly at a chromosomal DSB site (Aplan, 2006; Nussenzweig and Nussenzweig, 2010).

Indel-Seq reveals that most insertions are not generated from the direct ligation of extrachromosomal DNA fragments but require contact between the acceptor and donor loci followed by recombination-dependent events. First, we established that sites of interactions between the acceptor DSB on chromosome 17 and other AsiSI-dependent breaks along that chromosome correspond to donor sites for insertions (Fig. 2C). Clustering of DSBs within DNA repair domains is in part driven by nuclear actin polymerization involving the WASP-ARP2/3 pathway (Caridi et al., 2018; Schrank et al., 2018). These movements, which facilitate homology-dependent repair (Schrank and Gautier, 2019) but also enable pathologic translocations (Zagelbaum et al., 2022) are as well a driving force for genome insertions, another pathological form of structural variation and affect primarily Class (1) and (2) insertions (Fig. 2A-B). Interestingly, our recent study indicates that small insertions proximal to a translocation donor chromosome can be found at translocation junctions, suggesting that insertions could have a translocation-like intermediate (Zagelbaum et al., 2022). Finally, consistent with a requirement for contacts between chromosomes, Indel-Seq reveals that nearly 10% of insertions harbor more than one donor sequence. Note that the largest insert detected by Indel-Seq is 240 bp and that only mappable inserts > 20 bp are included in our analysis. Thus, we likely underestimate the proportion of multiple inserts, including those that generate inserts > 240bp. If the majority of insertions arose from independent “cut and paste” mechanisms, the probability of multiple insertions would decrease sharply as the number of fragments increases and become neglectable. Multiple insertions are compatible with sequential contacts between the acceptor loci and multiple loci on different chromosomes (Fig. 2F).

Our data further indicate that intra- and inter-chromosomal contacts trigger subsequent templated insertions. An insertion mechanism requiring single or multiple contact between chromosomes differs from the interpretation of studies in yeast (Yu et al., 2018) proposing that insertions in DNA2-deficient cells arose from fragmented chromosomes.

### Recombination mechanisms drive insertions downstream of chromosome contacts

Nuclear actin driven movements affect primarily resected DNA ends destined to be repaired by homology-directed repair (HDR). Therefore, mobile DNA lesions are already primed to undergo recombination-dependent repair (Schrank and Gautier, 2019). Consistent with that idea, we find that inhibiting DNA end-resection, the initial step of HDR (Symington and Gautier, 2011), alters the distribution of donor sequences derived from resected DNA tracks at cleaved AsiSI. We document a significant increase in insertions derived from the most AsiSI proximal sequences when MRE11 was inhibited (0 – 250bp) and an increase in more distal insertions (250 – 750bp) when BLM was inhibited. This suggests that resected ends, either at the TRIM37 acceptor site, at donor sites or both, can serve as template for insertions following microhomology-mediated base pairing, which is supported by the presence of short microhomologies at most insertion junctions (Fig. 3J). Moreover, reinforcing the idea that insertions are driven by recombination mechanisms, we find that chromatin-bound RAD51 is enriched at AsiSI donor loci. The top 20% of RAD51 enriched AsiSI loci contribute to nearly 50% of insertions. Finally, increasing RAD51 stability on chromatin by PARI down-regulation, a functional homolog of yeast Srs2 antirecombinase (Moldovan et al., 2012), significantly increases the numbers of insertions, those with multiple donor sequences being most affected, strongly suggesting that RAD51 filaments drive insertions.

We propose that resected DNA ends can also be processed by fill-in or endonucleolytic cleavage, yielding blunt ends compatible with subsequent NHEJ. Shieldin-CST-**Polα** complex binds to **ssDNA overhangs and facilitate 5’** fill-in (Mirman et al., 2018; Schimmel et al., 2021). Alternatively, ssDNA overhangs can undergo endonucleolytic processing to promote NHEJ, suggesting an additional mechanism by which DSBs are redirected to the NHEJ pathway when they fail to properly engage into HDR following DNA end-resection (He and Chowdhury, 2021; Zhao et al., 2021).

### Transcription drives genome insertions

Transcription is intimately linked to the DNA damage response and genome instability (Crossley et al., 2019; Petermann et al., 2022). Active transcription is a sensor of DNA lesions that can activate the DDR and subsequent repair. Transcription is also a significant cause of genome instability, as DNA unwinding generates fragile and exposed DNA intermediates. In addition, DNA polymerases are more processive than RNA polymerases; co-directional as well as head-to-head collisions are frequent yielding DSBs (Hamperl et al., 2017; Stork et al., 2016). Finally, DNA damage, DSBs in particular, stimulate transcription in the vicinity of DSBs (Cohen et al., 2018; Francia et al., 2012; Liu et al., 2021), as well as the formation of RNA-DNA hybrids (Bader and Bushell, 2020; Li et al., 2016; Michelini et al., 2017; Ohle et al., 2016). Multiple mechanisms are thought to yield RNA-DNA hybrid intermediates at DSBs, such as ongoing transcription pauses (Iannelli et al., 2017; Pankotai et al., 2012; Shanbhag et al., 2010), transcribed RNA fragment recruitment (Keskin et al., 2014; Welty et al., 2018), or de novo RNA synthesis initiation (Francia et al., 2012; Liu et al., 2021; Michelini et al., 2017). Whereas it is not entirely clear where RNA-DNA hybrid intermediates arise relative to the DSBs, RNA-DNA hybrids facilitate DNA end-resection during HDR (Crossley et al., 2019; Marnef and Legube, 2021).

R-loops are a documented source of genome instability that can be converted into DSBs following collision between DNA and RNA polymerases (Hamperl et al., 2017; Stork et al., 2016). We find that insertions originate preferentially around AsiSI sites that are active transcription loci. In cases where the AsiSI break is near a TSS, insertions from the transcribed side of the locus are enriched. Importantly, we show that inhibition of transcription by low-dose alpha-amanitin inhibits the formation of insertions, mostly Class 1 and Class 2. Consistent with the idea that RNA-DNA hybrids could be pathogenic intermediates in the generation of insertions, we demonstrate that R-loop resolution following RNASE H1 overexpression inhibits insertion formation, most significantly Class 2, which arise from R-loop rich regions (Fig. 3). Conversely, increasing transcription at telomeres following FANCM down-regulation resulted in increased telomeric insertions. All telomeres are distal from AsiSI breaks, yet, telomere insertion requires their own transcription. Furthermore, both Class 2 and telomere insertion depend on BLM/POLD3 mediated pathways (Fig 3). This suggests that BLM/POLD3-dependent break-induced replication (BIR) drives insertions, as observed during ALT telomere synthesis. Indeed, BLM is recruited to DSBs in transcriptionally active loci in response to the accumulation of R-loops, a process involving POLD3-dependent transcription-coupled DSB repair (Cohen et al., 2022). BIR involvement is consistent with the idea that microhomology-mediated BIR (MMBIR) participates in the generation of rearrangements in the human genome (Hastings et al. 2009). Our studies, however, uncover a transcription-coupled BIR mechanism distinct from MMBIR, thought to initiate at faulty replication forks.

Altogether this establishes that transcription drives genomic insertions. DSBs are thought to play a physiological role in transcription (Vitelli et al., 2017), therefore our findings could explain the generation of mutagenic genome insertions at transcription loci. Importantly, such rearrangements are predicted to occur in non-dividing cells, since they do not require active DNA replication. Transcription-induced rearrangements could explain age-dependent genome instability observed in post-mitotic neurons (Lu et al., 2004).

In summary, our data reveal key steps in the generation of potentially pathogenic insertions, leading to the comprehensive model proposed in Fig. 5C. Chromosome contacts between the rearranged loci are critical, in particular for donor sequences arising at the vicinity of DSBs: Class (1) and (2) insertions. We propose that contacts between the resected DNA end of the acceptor locus facilitate annealing and/or invasion of donor loci within HDR domains. Annealing is mediated by microhomologies (MHs) at donor loci resected DNA ends (1) or at the displaced DNA strand at donor R-loops loci (2). Our data is also consistent with subsequent BIR event(s), displacement of the invading strand and reassociation and processing of the templated donor with the acceptor locus followed by NHEJ. This model also applies for donor sequences arising spontaneously as DNA transactions intermediates (3), such as replication forks or ssDNA gaps.

## Methods

### Cell lines

U2OS and 293T cells were purchased from the American Type Culture Collection (ATCC). U2OS and 293T cells were maintained in Dulbecco’s Modified Eagle Medium (DMEM) supplemented with 10% fetal bovine serum. U2OS+ER-AsiSI (DivA) cells were obtained from Dr. Gaëlle Legube, and an early passage of DivA was used. DivA cells were maintained in DMEM supplemented with 10% fetal bovine serum supplemented with 1 μg/ml puromycin.

### Generation of knockdown and ectopic cell lines

#### Cas9 expression in U2OS cells

Lentiviruses were generated by transfection of pLenti-blast based vectors for Cas9 WT, Cas9 D10A, and Cas9 H840A together with pMD2.G and psPAX2 vectors into 293T cells using Lipofectamine 2000 (Invitrogen) according to the manufacturer’s protocol. The medium was replaced after 24 hours with fresh medium, and cells were maintained for an additional 24 hours. Supernatants containing virus were collected, passed through a 0.45 μm syringe filter, and concentrated with 4x virus concentrator solution (phosphate-buffered saline (PBS) at pH 7.2 containing 1.2 M sodium chloride, 40% [v/w] PEG-8000). Concentrated virus pellets were used to infect U2OS, and DivA cells supplemented with 0.5 μg/ml polybrene to enhance viral transduction. Cells were selected under 5 μg/ml Blasticidin-S (Invivogen) for two weeks.

#### gRNA expression in Cas9 expressing U2OS and DivA cells

Lentiviruses were generated by transfection of pKLV2.2-h7SKgRNA5(SapI)-hU6gRNA5(BbsI)-PGKpuroBFP-W (addgene #72666) based vectors for gRNAs targeting TRIM37 locus together with pMD2.G and psPAX2 vectors into 293T cells using Lipofectamine 2000 according to the manufacturer’s protocol. Virus-containing supernatants were harvested as described above, and then used to infect the Cas9 expressing cells described above. U2OS cells were selected under 2 μg/ml puromycin (Invivogen) for 3 days.

#### POLD3 shRNA-mediated knockdown

Lentiviruses were generated by transfection of pLK0.1-blast (Addgene #26655) based vectors for shRNA targeting human POLD3 (shPOLD3) or control vector together with pMD2.G and psPAX into 293T cells using Lipofectamine 2000 according to the manufacturer’s protocol. The virus infection was conducted as described above. Cells were selected under 5 μg/ml Blasticidin-S for two weeks.

#### Cell treatments

For treatment with 4-hyroxy-tamoxifen (4-OHT) (500 ng/ml), DivA cells were grown to ~70% confluence and then treated with 4-OHT for 48 or 72 hours. For all treatments: 4-OHT, NU7441 (20 μM), Mirin (20 μM), CK-666 (25 μM), or α-amanitin (1 μg/ml) of fresh medium and drug was added every 24 hours.

### Molecular cloning to generate knockdown and expression vectors

#### Mammalian expression vectors

cDNA encoding RNaseH1-GFP was amplified from RED (Addgene # 139835) vector using the primers below. RNaseH1 mitochondrial targeting sequence (MTS) was not amplified. Amplicons were cloned into XhoI- and AscI-digested pCS2 (+) FA vector. The pCS2 RNaseH1-GFP vectors were used for transfection in DivA cells using Neon transfection system (Invitrogen) according to manufacturer’s protocol. cDNA expression and transfection efficiency were assessed by GFP signal.

Forward XhoI-RNaseH1(M27): ACTCTCGAGGCCACCatgttctatgccgtgaggaggggccgcaag

Reverse AscI-eGFP (dCasN): ACTGGCGCGCCagctaagcctattgagtatttcttatccat

#### Lentiviral expression vectors

pLentiCas9-Blast (Addgene #52962) and pLentiCas9n(D10A)-Blast (Addgene #635993) were purchased from Addgene. To generate pLentiCas9 (H840A)-Blast vector, cDNAs encoding Cas9 protein were amplified from pLentiCas9-Blast vector using the primers below. Amplicons were cloned into HindIII- and BamHI-digested pCS2 (+) FA vector. pCS2 Cas9 vectors was used for Site-directed mutagenesis to generate pCS2 Cas9 H840A vector. Mutagenic primers were used for amplification, followed by KLD reactions (New England Biolab) according to the manufacturer’s protocol. Cas9 H840A cDNAs were amplified from pCS2 Cas9 H840A and cloned into AgeI- and BamHI-digested pLentiCas9-Blast vector to generate the pLentiCas9 (H840A)-Blast vector.

Forward HindIIIAgeI Cas9: ACTAAGCTTaccggttctagagcgctgccaccatggac

Reverse BamHI Cas9: ACTGGATCCttatcgtcatcgtctttgtaatc

Forward Cas9 H840A: gatgtggacGCtatcgtgcctcagagctttc

Reverse Cas9 H840A: gtagtcggacagccggttgatgtcc

#### shRNA vectors

pLK0.1 based vectors for the expression of shRNAs targeting POLD3 gene was generated by cloning the double stranded oligonucleotides listed below into EcoRI- and AgeI-digested pLK0.1-Blast vector.

Sense shPOLD3: CCGGCAATTAGTGGTTAGGGAAAAGCTCGAGCTTTTCCCTAACCACTAATTGTTTTTG

Anti-sense shPOLD3: AATTCAAAAACAATTAGTGGTTAGGGAAAAGCTCGAGCTTTTCCCTAACCACTAATTG

The target sequence is CAATTAGTGGTTAGGGAAAAG

#### sgRNA vectors

pKLV2.2-h7SKgRNA5(SapI)-hU6gRNA5(BbsI)-PGKpuroBFP-W based vector for gRNAs targeting TRIM37 locus expression was generated by cloning the double stranded oligonucleotides listed below into BbsI- and SapI-digested vector sequentially.

Sense hU6 BbsI TRIM37 (+7): CACCGgccccgcaacgcgggaact

Antisense hU6 BbsI TRIM37 (+7): AAACagttcccgcgttgcggggcC

Sense 7SK SapI TRIM37 (+53): CTCGcttggcgactcgctgcctc

Antisense 7SK SapI TRIM37 (+53): AACgaggcagcgagtcgccaagC

#### siRNA-mediated knockdown in DivA cells

siRNA oligos were synthesized from Sigma-aldrich (10 nmol) and purified by desalting. To ensure the stability of siRNA, dTdT overhangs were added. siRNAs were transfected into DivA cells at a final concentration of 10 nM using lipofectamine 2000 according to the manufacturer’s protocol. After 24 hours, 4-OHT was added for the experimental purpose.

siRNA sequences:

siPARI: AGGACACAUGUAAAGGGAUUGUCUA

siBLM: AUCAGCUAGAGGCGAUCAA

siFANCM: AAGCUCAUAAAGCUCUCGGAA

siFANCM-U: CCGGAUGAGUGAACAAUACUU

siLuc: UCGAAGUAUUCCGCGUACGUU

### Indel-seq procedure

#### Preparation of genomic DNA and *in vitro* AsiSI digestion

Genomic DNA was prepared using Puregene core kit A (Qiagen) according to the manufacturer’s protocol. DNA quality was checked by electrophoresis on 1% agarose gel. For in vitro AsiSI digestion, ten micrograms of genomic DNA were digested by 100 unit of AsiSI enzyme at 37 °C for 16 hours. Digested DNA was purified by ethanol precipitation for 16 hours at −20 °C. Precipitated DNA was washed twice with 70% ethanol, air dried, and eluted into water.

#### Library preparation for Indel-seq

Two hundred nanogram of DNAs were used as a template for polymerase chain reaction (PCR) by using EmeraldAmp MAX PCR mater mix (Takara). First 30 cycles of PCR were performed using forward and reverse primers below, followed by 14 PCT cycles with P5I5- and P7I7-primers. In each step, at least eight separate tubes were used for PCR and then pooled and purified by using PCR and gel extraction kit (Macherey-Nagel) according to manufacturer’s protocol. The quality and concentration of purified amplicons were checked by Bioanalyzer High Sensitivity DNA ChIP (Agilent) and Qubit dsDNA HS assay Kit (Invitrogen) at Herbert Irving Comprehensive Cancer Center Shared Resource.

#### PCR primer sequences

Forward TRIM37: AATTCGCAAACACCAACCGT

Reverse TRIM37: TCTGAAGTCTGCGCTTTCCA

Forward P5-I5-NNNNNN-barcode-TRIM37:

AATGATACGGCGACCACCGAGATCTACACTCTTTCCCTACACGACGCTCTTCCGATCTNNN NNNbarcodeAATTCGCAAACACCAACCGT

Reverse P7-I7-NNNNNN-TRIM37: CAAGCAGAAGACGGCATACGAGATCGGTCTCGGCATTCCTGCTGAACCGCTCTTCCGATCTNNNNNNTCTGAAGTCTGCGCTTTCCA

#### MiSeq

MiSeq was performed by using Illumina MiSeq Reagent Kit v3 (600-cycle) according to manufacturer’s protocol “Denature and Dilute Libraries Guide”. Briefly, four nanomole of pooled library was denatured by adding 0.2 N Sodium hydroxide solution and diluted to 20 pM. 12 pM library was mixed with 50% PhiX for better sequencing results. FASTAQ only was used as a run setting. TruSeq Nano DNA was used as a Library Prep workflow. TruSeq DNA Single Indexes (A,B), 0 Index reads, and Adapter trimming/read2 were set. Over the course of 72 hours, 301×2 paired end reads were performed.

#### INDEL-seq analysis

FASTAQ file demultiplexing was performed using fastq-multx program. Then, paired-end reads were merged into full sequence segments using PEAR based on read overlaps (Zhang et al., 2014). Lastly, the reads were aligned against the reference sequence to generate SAM file. Insertions and deletions were identified by parsing the CIGAR string.

#### Donor sequence analysis

Insertion sequences >20bp were aligned to human reference genome (hg19) with the BLAT program. Genomic regions obtained from BLAT were then annotated using R package ChIPseeker (Yu et al., 2015). The distance between the donor sequence and the nearest AsiSI motif was calculate for 1211 AsiSI motifs (GCGATCGC) in the human genome. For promoter-proximal DSBs analysis, USP22 (Chr17: 20946300), TRIM37 (Chr17: 57184296), CSNK1D (Chr17: 80231110), ASXL1 (Chr20: 30946312), KDELR3 (Chr22: 38864101) loci were used for the analysis. Microsoft Excel and GraphPad Prism 8 were used to process the generated sequence data.

#### ChIP-seq data analysis

ChIP-seq reads were aligned to the hg19 genome assembly using bwa mem. Peaks were called using macs2 with the default setting. Following ChIP-seq and BLESS data set was used: S9.6 ChIP (ERR2224138, ERR2224139) (Cohen et al., 2018), BLESS (ERR2008259, ERR2008260) (Clouaire et al., 2018).

#### Hi-C

Chromosome conformation capture experiments were performed as previously described33 with some modifications. Briefly, 5 million DivA cells/library were crosslinked with 1% formaldehyde and lysed. After digesting chromatin with 400 units of DpnII overnight, DNA ends were labeled with biotinylated dATP using 50 units Klenow DNA polymerase. Blunt-end ligation was performed with 50 units T4 Ligase at 16°C for 4 hours, followed by reverse crosslinking with 400 mg/ml proteinase k at 65°C overnight. DNA was purified using phenol/chloroform extraction and ethanol precipitation, and concentrated on a 30 kDa Amicon Ultra column. Biotin was removed from unligated ends in 50ml reactions using 50 units T4 DNA polymerase/5 mg DNA. Following DNA sonication (Covaris S220) and Ampure XP size fractionation to generate DNA fragments of 100-300 bp, DNA ends were repaired using 7.5U T4 DNA polymerase, 25U T4 polynucleotide kinase, and 2.5 U Klenow DNA polymerase. Libraries were enriched for ligation products by biotin pulldown with MyOne streptavidin C1 beads. To prepare for sequencing, A-tailing was performed using 15 units of Klenow DNA polymerase (3’-5’ exo-) and Illumina TruSeq DNA LT kit Set A indexed adapters were ligated. Libraries were amplified in PCR reactions for 5-7 cycles and subjected to Ampure XP size selection prior to sequencing on an Illumina HiSeq 4000 machine using the Paired End 50 bp module. Two biological replicates were performed.

#### Hi-C Analysis

Paired-end 50bp reads were processed using the distiller pipeline(Goloborodko et al., 2019). First, reads were mapped to hg19 reference genome, using BWA-MEM in single sided mode (- SP). Alignments were then parsed, classified, and filtered using pairtools(Goloborodko et al., 2019). The resulting valid pairs included uniquely mapped and rescued pairs with a minimum mapping quality of 30. Valid pairs were aggregated into binned contact matrices and kept as multi-resolution cooler files(Abdennur and Mirny, 2020) for subsequent analyses. Paired reads from replicate libraries were pooled prior to filtering for PCR duplicates. All Hi-C contact matrices were normalized by iterative correction(Imakaev et al., 2012), excluding the first 2 diagonals. Downstream analyses were performed using cooltools version 0.3.2(Venev et al., 2020) and python 3.7.10.

#### PCAWG indel analysis

We downloaded the indel calling result of PCAWG study (in total 2,778 WGS samples). Low quality indel candidates were filtered out. Gene strand and Transcription Start Site (TSS) information were extracted from Ensembl genome browser. Within each cancer type, for each gene, the total numbers of indel candidates inside the upstream and downstream 50kb windows from TSS were compared searching for potential bias using prop.test in R.

The custom scripts used in the data analysis can be find at: https://github.com/RabadanLab/INDEL-seq

#### Sanger sequencing of inserts

TRIM37 amplicons containing insertions were purified using gel extraction kit (Macherey-Nagel) after 1.5 % agarose gel gel-electrophoresis, and then cloned into pGEMT-easy vector (Promega) according to manufacturer’s protocol. further plasmid DNAs of each clone were prepared by using Nucleospin Plasmid (Macherey-Nagel) and sent for Sanger sequencing (Azenta) using T7 and SP6 sequencing primers. Over 500 sanger sequencing result, 162 clones harboring unique insertion sequences were selected for the analysis.

#### Southern blot

Southern blot was performed as previously described (Lai et al., 2017), TRIM37 amplicons were run into 1% agarose gel in 1× tris-acetate-ethylenediaminetetraacetic acid (TAE) buffer for 2 hours with 3 V/cm. The gel (SYBR DNA) was imaged using an Accuris blue light Smartdoc (Sigmaaldrich). This is followed by depurination with 0.2 M hydrochloride for 15 min, denaturation with 0.5 M sodium hydroxide and 1.5 M sodium chloride for 20 min, and neutralization with 0.5 M Tris-HCl buffer (pH 8.0) and 2 M sodium chloride for 20 min. Gels were transferred onto a positively charged nylon membrane using VacuGene XL (GE Healthcare) and cross-linked in a UV Stratalinker. The membrane was then hybridized with DIG-labeled telomere probe in 42 °C 16 hours. Hybridized membrane was washed in 2 X Saline Sodium Citrate (SSC) and 0.1% (v/w) Sodium Dodecyl Sulfate (SDS) twice for 15 minutes, followed by another wash in 0.5 X SSC and 0.1% SDS twice for 15 minutes. The membrane was transferred to a blocking buffer (1 M maleic acid solution pH7.4, 1% nucleotide blocking reagent (Roche)) for 30 minutes and then incubated in antibody-solution (anti-dioxigenin-AP fragments in blocking buffer). The membrane was washed with Wash buffer (1M maleic acid solution pH7.4, 0.3% tween 20), followed by chemiluminescent reaction in CDP-star solution (Roche). The membrane was sealed in film and imaged using a Li-Cor Odyssey imager.

#### RNA extraction and Northern blot

Total RNA was extracted by using RNeasy mini kit (Qiagen) according to the manufacturer’s protocol. 200 ng of total RNA were run in 1% agarose gel in 1X TAE buffer for 1 hours with 2 V/cm. The gel staining was imaged for total RNA. To detect the telomeric RNA (TERRA), 1 μg of total RNA were slot blotted, followed by hybridization with telomere C-probe and imaged as described in southern blot.

#### Cell lysate extraction and Western blot

Western blot was performed as previously described (Li et al., 2022), cells were incubated with lysis buffer (50 mM HEPES pH 7.4, 150 mM NaCl, 10% glycerol, 1% Triton X-100, 1 mM EDTA, 100 mM sodium fluoride, 2mM sodium orthovanadate, 20 mM sodium pyrophosphate, 0.5 mM dithiothreitol, 2 mM PMSF) supplemented with cOmplete Protease Inhibitor Cocktail (Roche) and PhosSTOP (Sigma) on ice for 1 hour. Protein concentrations of cell lysates were measured after centrifugation at 20,817g for 10 minutes at 4 °C using a Micro BCA Protein Assay Kit (Invitrogen). 50 μg of total proteins were analyzed by SDS-PAGE and Western blotting. The following antibodies were purchased from the indicated commercial sources: anti-BLM (1:1000; Bethyl laboratories A300-110A), anti-POLD3 (1:500; Proteintech, 21935-a-AP), anti-β-Actin (1:1000; C-4, Santa Cruz Biotechnology, sc-47778). anti-rabbit immunoglobulin G (IgG) (H + L) (Dylight 800 conjugates) and anti-mouse IgG (H + L) (Dylight 680 conjugates) (Cell Signaling) were used as secondary antibodies.

#### Quantification and Statistical Analysis

Statistical analyses for the genomic experiments were performed using standard genomic statistical tests as described above and figure legends. Statistical analysis for the Indel-seq experiments were performed using GraphPad Prism 8. All tests and p values are provided in the corresponding figures and figure legends.

## Acknowledge

We thank Molecular Pathology Shared Resources of the Herbert Irving Comprehensive Cancer Center (HICCC) at Columbia University for the bioanalyzer analysis. We also thank Eunhee Choi at Columbia University for the Western blot analysis. This work was supported by the following NIH/NCI/NHGRI grants: CA245259 (J.M.), CA174653 (J.G., R.R., and Ju.Z), F30-CA250166 (Je.Z.), CA197606 (J.G.), and HG003143 (J.D.). J.D. is a HHMI investigator.

