## Supplemental Figures for "Mechanisms of insertions at a DNA double-strand break"

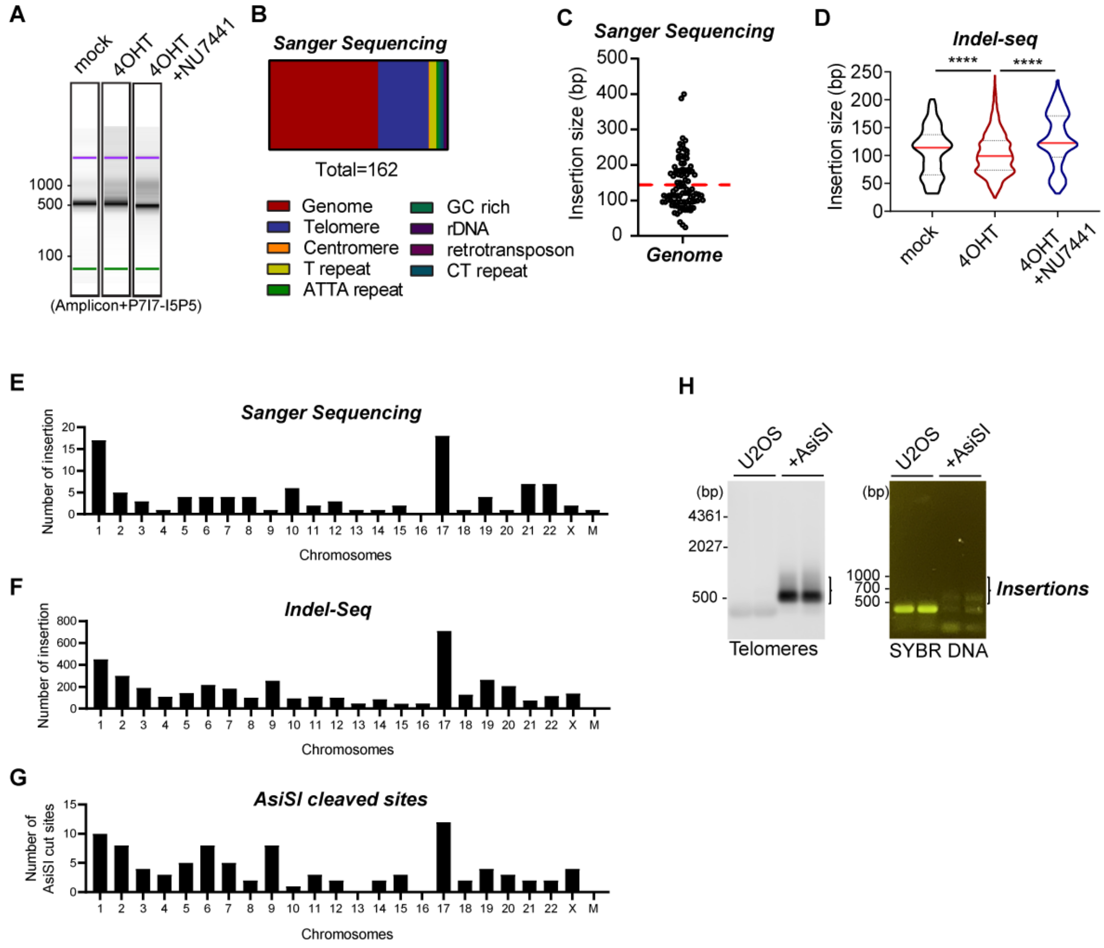


**Figure S1, related to Figure 1**

(A) Examples of bioanalyzer profiles of PCR amplicons spanning the AsiSI site at the TRIM37 locus in control (mock), 4OHT treated, and NU7441 and 4OHT treated DIvA cells. The amplicons were tagged with P5I5 and I7P7 sequences for MiSeq run. (B) Origins of donor sequences from Sanger sequencing. (C) Distribution of insertion size from Sanger sequencing. (D) Insertion size shown as violin plots in control (mock), 4OHT treated, and NU7441 treated DIvA cells. Red lines indicate median. (E-F) Chromosomal origin of donor sequences from Sanger sequencing (E) and Indel-seq (F). (G) Chromosomal distribution of AsiSI frequently cut site. (H) Telomere Southern blot (TTAGGG) (left) and SYBR DNA staining (right) of amplicons spanning the AsiSI site at the TRIM37 locus in U2OS and U2OS ER-AsiSI+4OHT (+AsiSI) cells.

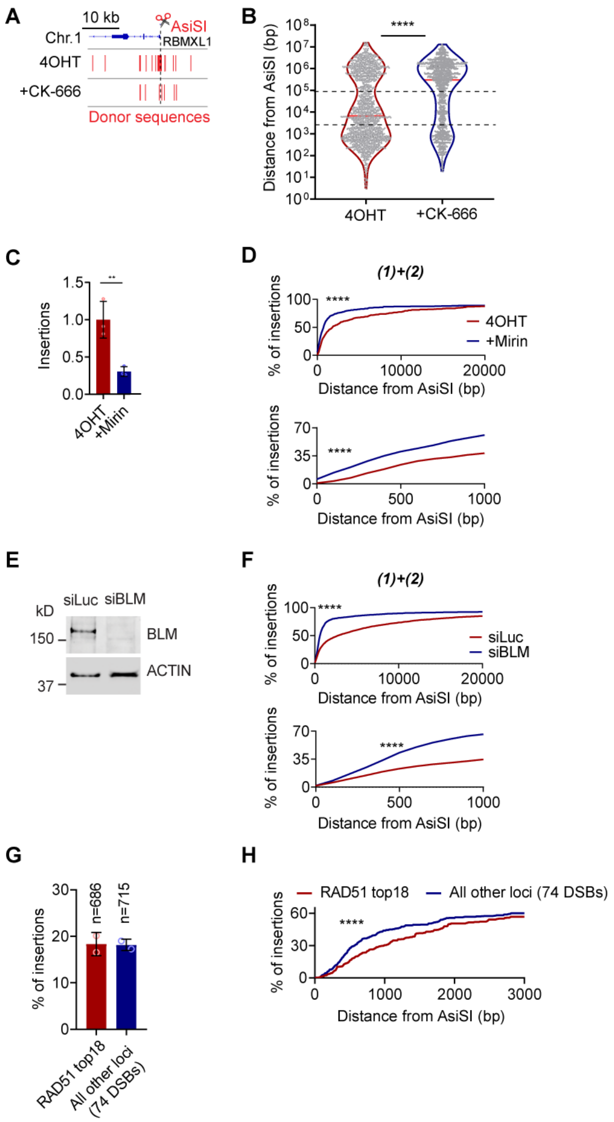


**Figure S2, related to Figure 2**

(A) Genome browser view of RBMXL1 locus (top); distribution of donor sequences in 4OHT treated (middle) or 4OHT and 50 μM CK-666 treated DIvA cells (bottom). (B) Distribution of donor sequences as a function of the distance to the nearest AsiSI site in 4OHT +/- CK-666 treated DIvA cells. Bar graphs represent median. (C) Relative number of insertion events in 4OHT +/- Mirin treated cells. Columns are normalized to the frequency of insertions in cells treated with 4OHT. Mean and standard deviation. (D) Cumulative frequency of the distribution of donor sequences proximal to DSBs ((1)+(2); < 100 kb)), as a function of the distance from the nearest AsiSI site in 4OHT +/- Mirin treated cells. *P* calculated with Kolmogorov-Smirnov test. (E) BLM protein expression in DIvA cells following siRNA knockdown of the BLM gene. (F) Cumulative frequency of the distribution of donor sequences proximal to DSBs ((1)+(2); < 100kb)), as a function of the distance from the nearest AsiSI site in control and BLM-depleted cells. *P* calculated with Kolmogorov-Smirnov test. (G) Percentage of donor sequences which are derived from 18 most RAD51-enriched DSBs (RAD51 top 18) and for all other loci (74 DSBs). (H) Cumulative frequency of the distribution of donor sequences derived from RAD51-enriched DSBs (top 18) and all other loci, as a function of the distance from the nearest AsiSI cut site. *P* calculated with Kolmogorov-Smirnov test.

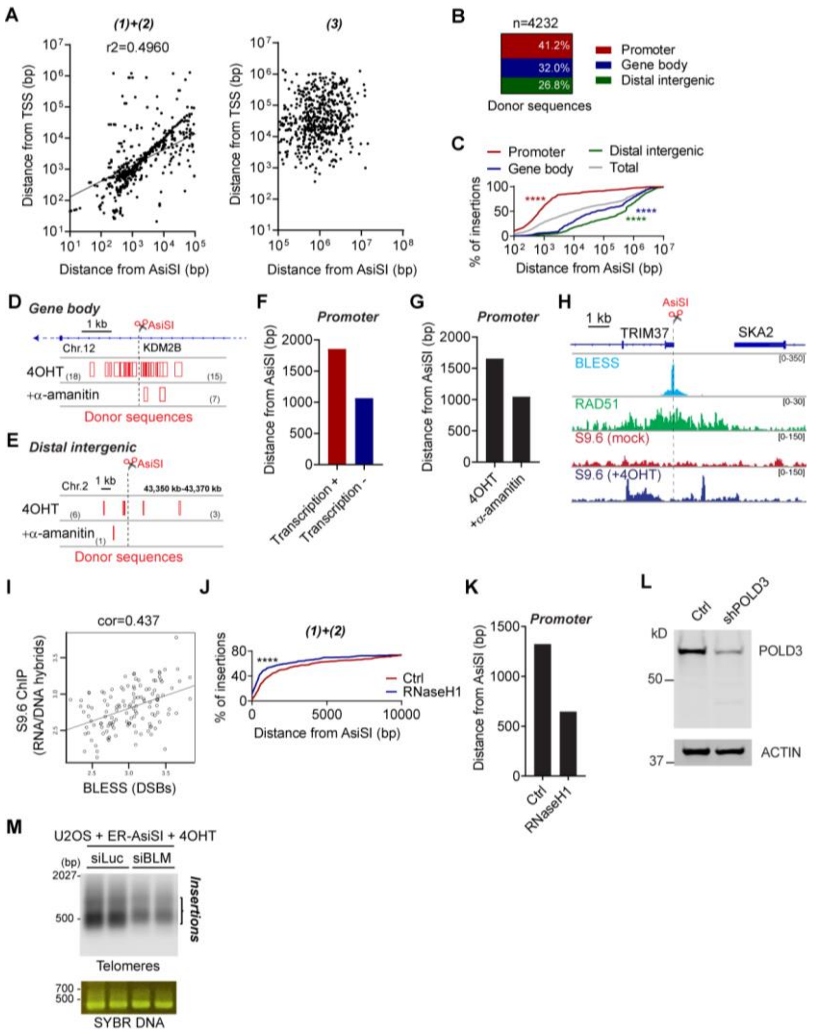


**Figure S3, related to Figure 3**

(A) Log-log correlation between the distance from the nearest AsiSI site versus the distance from TSS for donor sequences which are proximal to AsiSI site (Class 1 and 2, left) and distal to AsiSI site (Class 3, right). The correlation coefficient (R-squared) is shown. (B) Distribution of donor sequences originating from promoter, gene body, and distal intergenic regions. (C) Cumulative frequency of the distribution of donor sequences originating from promoter, gene body and distal intergenic regions as a function of the distance from the nearest AsiSI site. *P* calculated with Kolmogorov-Smirnov test. (D-E) Genome browser view of KDM2B locus (gene body) (D) and of an intergenic locus (E). The distribution of donor sequences following 4OHT treatment +/- alpha-amanitin are shown below. (F-G) Quantification of the distance of donor sequences (derived from AsiSI nearby TSS) from AsiSI sites in transcribed and non-transcribed regions (F), and in DIvA cells treated with 4OHT +/- $\alpha$-amanitin (1 $\mu$g/ml) (G). (H) Genome browser view of the TRIM37 locus (top), BLESS, RAD51 and S9.6 ChIP-seq, and the distribution of donor sequences. (I) Log-log correlation between BLESS signal and S9.6 ChIP-seq for the 128 most frequently cut AsiSI sites in DIvA cells. Pearson correlation ("r") is shown. (J) Cumulative frequency of the distribution of donor sequences proximal to DSBs ((1)+(2); < 100 kb)), as a function of the distance from the nearest AsiSI site in control and RNaseH1 cDNA overexpressing DIvA cells. *P* calculated with Kolmogorov-Smirnov test. (K) Quantification of the distance of donor sequences (derived from AsiSI nearby TSS) from AsiSI sites in control and RNaseH1 cDNA overexpressing DIvA cells. (L) POLD3 protein expression in DIvA cells following shRNA knockdown of the POLD3 gene. (M) Telomere Southern blot (TTAGGG) (top) and SYBR DNA staining (bottom) of amplicons spanning the AsiSI site at the TRIM37 locus in control (siLuc) and BLM-depleted DIvA cells.

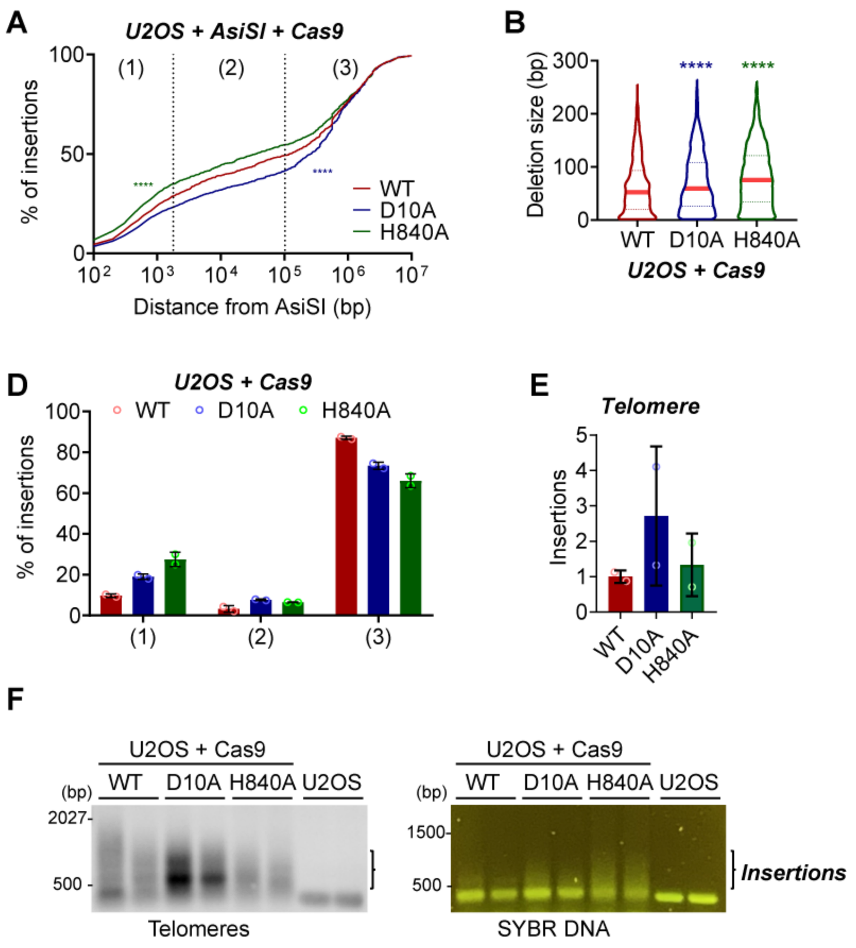


**Figure S4, related to Figure 4**

(A) Cumulative frequency of the distribution of donor sequences in 4OHT-treated DivA cells expressing WT and nickases Cas9 and gRNAs targeting TRIM37 locus as a function of the distance from the nearest AsiSI site. *P* calculated by Kolmogorov-Smirnov test. (B) Deletion sizes in U2OS cells expressing WT and nickases Cas9 and gRNAs targeting TRIM37 locus. *P* calculated with Mann-Whitney test. (C) Number of complex insertion events in U2OS cells expressing WT Cas9 and nickases Cas9 and gRNAs targeting TRIM37 locus. (D) Frequency of donor sequences from each class in U2OS cells expressing WT and nickases Cas9 and gRNAs targeting TRIM37 locus. (E) Relative number of telomere insertion events in U2OS cells expressing WT and nickases Cas9 and gRNAs targeting TRIM37 locus. Columns are normalized to the frequency of insertions in Cas9 WT. Mean and standard deviation. (F) Telomere Southern blot (TTAGGG) (left) and SYBR DNA staining (right) of amplicons spanning the Cas9 break site at the TRIM37 locus in U2OS cells expressing WT Cas9 and nickases Cas9 and gRNAs targeting TRIM37 locus.

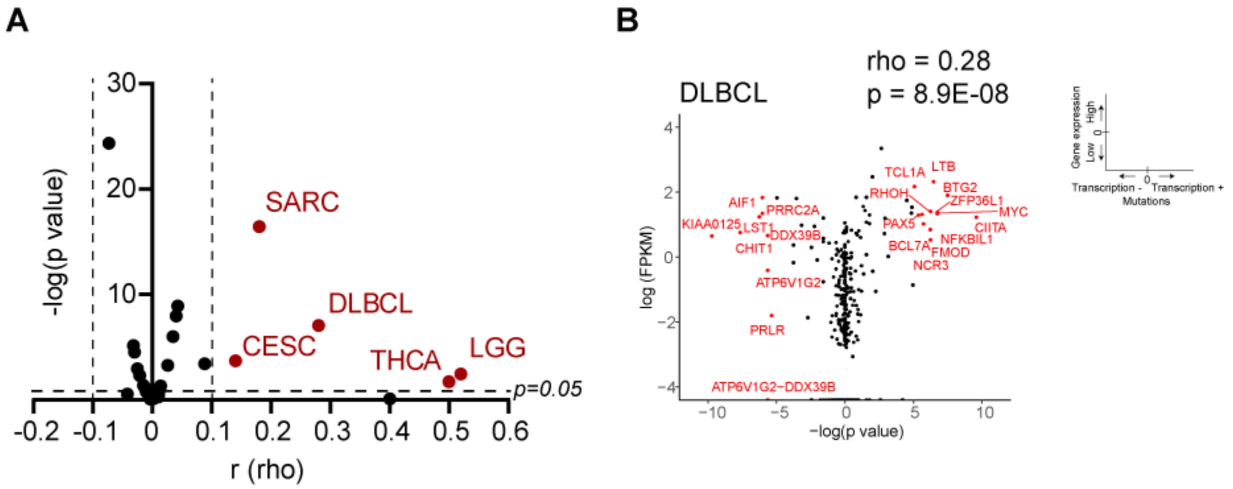


**Figure S5, related to Figure 5**

(A) Analysis of transcription-associated mutagenesis bias in 29 tumor types. x-axis: r (rho), correlation between transcription-associated bias and gene expression levels, y-axis: negative logarithm p value. r = ± 0.1, p=0.05: cut-off. (B) Analysis of transcription-associated bias in mutations in the diffuse large B-cell lymphoma (DLBC). x-axis: negative logarithm p value, positive: transcription-associated bias in transcribed side (Transcription +), negative: transcription-associated bias in un-transcribed side (Transcription -). y-axis: gene expression levels; logarithm of Fragments Per Kilobase of transcript per Million mapped reads (FPKM).
